# RAS mutation patterns arise from tissue-specific responses to distinct oncogenic signaling

**DOI:** 10.1101/2021.12.10.472098

**Authors:** Ozgun Erdogan, Nicole L.K. Pershing, Erin Kaltenbrun, Nicole J. Newman, Jeffrey I. Everitt, Christopher M. Counter

## Abstract

Despite multiple possible oncogenic mutations in the proto-oncogene *KRAS*, unique subsets of these mutations are detected in different cancer types. As KRAS mutations occur early, if not being initiating, these mutational biases are ostensibly a product of how normal cells respond to the encoded oncoprotein. Oncogenic mutations can impact not only the level of active oncoprotein, but also engagement with effectors and other proteins. To separate these two effects, we generated four novel inducible *Kras* alleles encoded by the biochemically distinct mutations G12D versus Q61R encoded by native (nat) rare versus common (com) codons to produce either low or high protein levels. Each allele induced a distinct transcriptional response in normal cells. At one end of the spectrum, the *Kras^natG12D^* allele induced transcriptional hallmarks suggestive of an expansion of multipotent cells, while at the other end, the *Kras^comQ61R^* allele exhibited all the hallmarks of oncogenic stress and inflammation. Further, this dramatic difference in the transcriptomes of normal cells appears to be a product of signaling differences due to increased protein expression as well as the specific mutation. To determine the impact of these distinct responses on RAS mutational patterning *in vivo*, all four alleles were globally activated, revealing that hematolymphopoietic lesions were sensitive to the level of active oncoprotein, squamous tumors were sensitive to the G12D mutant, while carcinomas were sensitive to both these features. Thus, we identify how specific KRAS mutations uniquely signal to promote the conversion of normal hematopoietic, epithelial, or squamous cells towards a tumorigenic state.

## Introduction

Why specific driver mutations track with different cancers is unknown, yet holds the key to understanding the origins of cancer, with implications for early detection and prevention. This is particularly well illustrated with the small GTPase KRAS. Single point mutations at one of three hotpot positions (G12, G13, and Q61) result in six possible substitutions that inhibit the intrinsic or extrinsic GTPase activity of the protein, rendering KRAS constitutively GTP-bound and active, which is well known to be oncogenic (***Simanshu et al., 2017***). Despite 18 possible oncogenic mutations, specific subsets of these mutations tend to be found in specific cancer types. For example, G12C is the most common KRAS mutation in non-small cell lung cancer while it is Q61H in plasma cell myeloma (***Li et al., 2018; Prior et al., 2020***). Mice similarly exhibit a bias of specific Kras mutations towards different cancer types, as different mutant *Kras* alleles have different tumorigenic potential when activated in different tissues (***Li et al., 2018; Poulin et al., 2019; Winters et al., 2017; Wong et al., 2020; Zafra et al., 2020***). While this tissue ‘tropism’ of cancers towards specific KRAS mutations has been appreciated for decades (***Bos, 1989***), the underlying mechanism is unclear.

Variation in the ability of specific KRAS mutants to be tumorigenic (or not) in different tissues ostensibly results from differences in oncogenic signaling between mutants. Generally speaking, oncogenic signaling is a product of the amplitude of the signal (quantitative signaling) and/or the effector pathway engaged (qualitative signaling). In terms of quantitative signaling, different mutations can exhibit different degrees of activation (GTP-loading) and/or different sensitivities to positive (Ras GTP exchange factors, RASGEFs) or negative (RAS GTPase activating proteins, RASGAPs) regulators (***Gebregiworgis et al., 2021; Lu et al., 2016; Muñoz-Maldonado et al., 2019; Simanshu et al., 2017***). Various methods to manipulate quantitative Kras signaling, such as through modulating recombination rates (***Singh et al., 2020***), homozygous expression of the mutant allele (***Burgess et al., 2017***), changing codon usage to increase translation (***Pershing et al., 2015***), additional pharmacologic activation of the mitogen activated protein kinase (MAPK) pathway (***Cicchini et al., 2017***), and so forth (***Li et al., 2018***), all affect tumorigenesis. On the other hand, perhaps one of the best examples of qualitative differences in KRAS signaling is the G12R mutant. Unlike the more canonical G12D mutation, the G12R mutant exhibits reduced binding the RAS effector PI3K*α* and PI3K/AKT signaling (***Hobbs et al., 2020; Zafra et al., 2020***). Activating an inducible *Kras^G12R^* allele in the pancreas led to early premalignant lesions compared to the much more tumorigenic *Kras^G12D^* allele (***Zafra et al., 2020***). Despite an appreciation that different KRAS mutations can manifest in quantitative or qualitative signaling differences, how these two signaling outputs impact the mutational patterns of this oncogene was unclear.

As KRAS mutations are often initiating, being sufficient to induce tumorigenesis in mice and truncal in many human cancers, the bias of specific mutations towards distinct cancers may arise from tissue-specific responses of normal cells to quantitative and qualitative features of KRAS signaling (***Li et al., 2018***). Determining the immediate response of normal cells to different Kras mutations *in vivo* thus holds the key to understanding the mutational patterning of this oncogene. Identifying a point mutation arising in *KRAS* from the cell-of-origin prior to becoming a tumor is challenging in humans. Mice, on the other hand, provide an ideal model system to experimentally explore this phenomenon, as the point of tumor initiation can be precisely defined using inducible oncogenic *Kras* alleles. To thus determine why specific KRAS mutations have such a strong bias for different cancer types we created four novel inducible murine *Kras* alleles designed with very different oncogenic mutations that were expressed at either low or high levels. In this way, tissues sensitive to quantitative signaling would develop tumors dependent on the activation status of the Kras oncoprotein, whereas those sensitive to qualitative signaling would develop tumors dependent upon the mutation type.

We chose two completely different oncogenic mutations for these experiments, namely G12D and Q61R. G12D places a negatively charged head group into the catalytic cleft of RAS and blocks extrinsic (RASGAP-mediated) GTPase activity (***Parker et al., 2018***). On the other hand, Q61R replaces the catalytic amino acid with one that has a positively charged headgroup, disrupting the position of the active site water molecules necessary for intrinsic GTP hydrolysis (***Buhrman et al., 2010***). Q61 is also essential for extrinsic GTP hydrolysis, as it stabilizes the transition state via hydrogen bonds to the *γ*-phosphate and nucleophilic water while providing another hydrogen bond to the GAP arginine finger (***Grigorenko et al., 2007; Kotting et al., 2008; Rabara et al., 2019; Scheffzek et al., 1997***). Comparing these two mutants directly reveals Q61R to have significantly lower GTP exchange and GTP hydrolysis rates than G12D (***Burd et al., 2014; Rabara et al., 2019***), akin to other substitutions at these two positions (***Gebregiworgis et al., 2021; Smith et al., 2013***). In those few cases in which the tumorigenic potential of the G12D and Q61R mutants have been directly compared in mice, tissue specific expression of Nras^G12D^ or Kras^G12D^ was less potent than their Q61R counterparts at inducing melanoma (***Burd et al., 2014***) or myeloproliferative neoplasm (***Kong et al., 2016***), respectively.

To parse out the contribution of quantitative oncogenic signaling, the *Kras* alleles encoding these two different mutations were expressed at different levels by manipulating their codon usage. Namely, the first three coding exons were fused and encoded by either their native rare codons, which is known to retard protein translation, or common codons to increase translation (***Lampson et al., 2013; Pershing et al., 2015***). We chose the novel approach of altering mammalian codon usage to modulate protein expression (***Pershing et al., 2015***), as no additional elements are required to change protein levels, providing a simple, reproducible, and uniform way of precisely controlling *Kras* levels in mice and derived cell lines.

These four alleles were activated and immediately thereafter the transcriptome of normal cells was determined, which revealed that each allele induced a specific transcriptional response in normal cells. Increased expression shifted the transcriptional hallmarks consistent with an expansion of multipotent cells to that of oncogenic stress and inflammation. Changing the mutation shifted the hallmark of estrogen response in the G12D mutant to that of the p53 pathway and DNA repair in the Q61R mutant. To determine how these two types of signaling contribute to RAS mutational patterning *in vivo*, all four alleles were globally activated, revealing that hematolymphopoietic lesions were sensitive to the level of active oncoprotein, squamous tumors were preferentially sensitive to the G12D mutant, while carcinomas tended to be sensitive to both these changes. Thus, we identify how specific KRAS mutations uniquely signal to promote the conversion of normal hematopoietic, epithelial, or squamous cells towards a tumorigenic state.

## Results

### A panel of *Kras* alleles designed to separate the effects of a mutation from the activity of the oncoprotein

To elucidate how specific cancers are driven by specific KRAS mutations, we reasoned that the contribution of quantitative signaling of the oncoprotein could be parsed out by simply changing the amount of protein made, while the contribution of qualitative signaling could be parsed out by using two different oncogenic mutants. To this end, we created the four novel *LSL-Kras^natG12D^*, *LSL-Kras^natQ61R^*, *LSL-Kras^comG12D^*, and *LSL-Kras^comQ61R^* alleles (***Figure 1A***) in which an LSL transcriptional/translational repressor sequence (STOP) flanked by *lox*P sites (***Jackson et al., 2001***) was engineered into the first intron of *Kras* after the non-coding exon 0 to provide temporal and special control of gene expression. This was followed by a fusion of coding exons 1 to 3 encoded by either their native (*nat*) rare codons, which are known to retard protein translation, or 93 of these rare codons converted to their common (*com*) counterparts to increase protein expression (***Figure 1—figure supplement 1,2***), and either a G12D or Q61R mutation. As noted above, each of these mutants alters RAS activity in a biochemically different manner (***Muñoz-Maldonado et al., 2019; Simanshu et al., 2017***), with Q61R reported to yield higher levels of active (GTP-bound) Ras (***Burd et al., 2014; Kong et al., 2016; Pershing et al., 2015***). This was followed by the next intron containing an FRT-NEO-FRT cassette for ES selection, which was excised via Flp-mediated recombination after which the Flp transgene was outbred. Finally, the remainder of the gene was left intact so as to generate the two Kras4a and Kras4b isoforms, as both contribute to tumorigenesis (***To et al., 2008***), potentially through unique protein interactions (***Amendola et al., 2019***).

**Figure 1.**
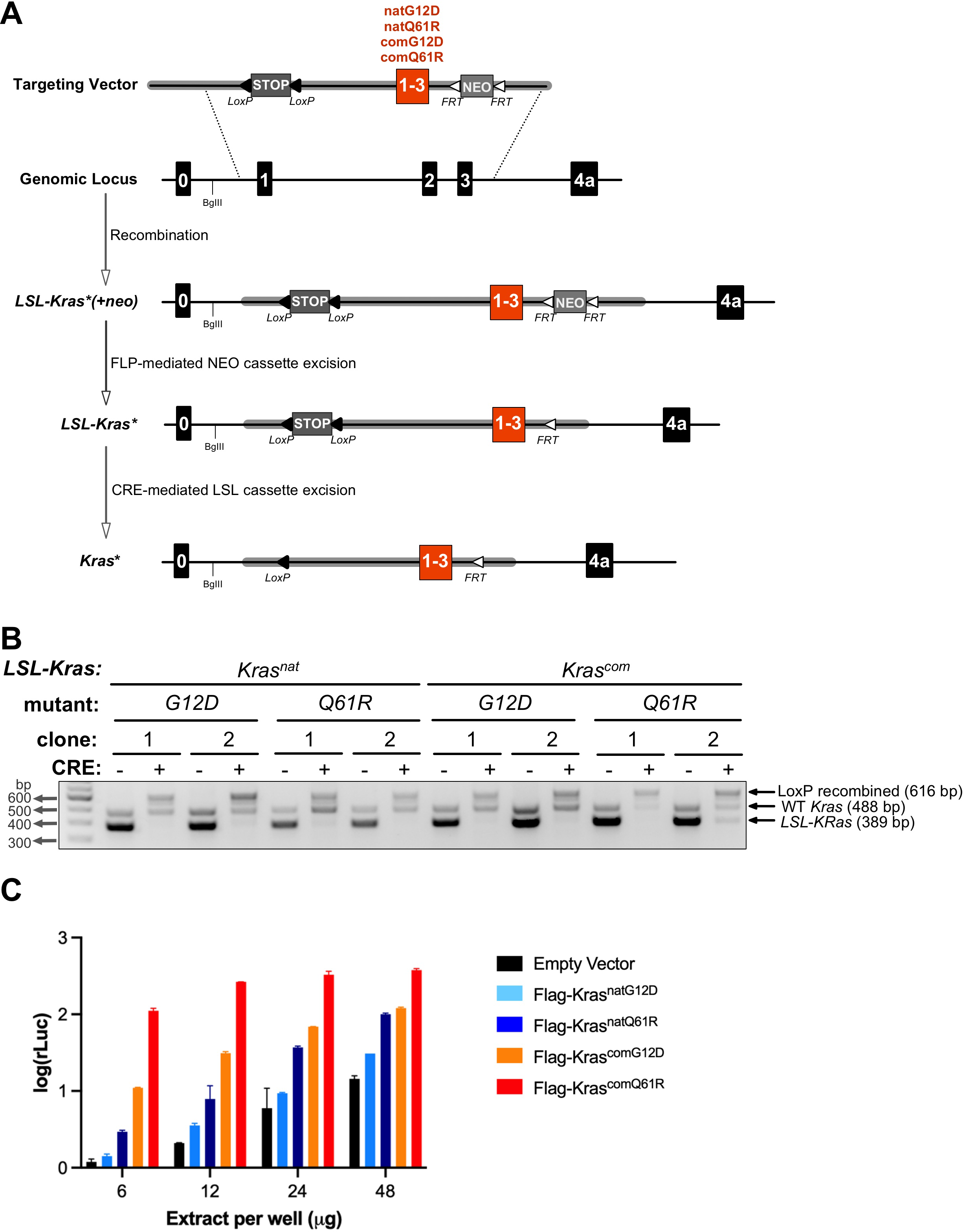
Conditional *LSL-Kras* alleles with different oncogenic mutations and codon usage. **(A)** Schematic of generating and activating *LSL-Kras* alleles with coding exons 1 to 3 encoded by native (*nat*) versus common (*com*) codons with either a G12D or Q61R mutation. **(B)** PCR genotyping of two independently derived MEF cultures (two biological replicates) with the indicated *LSL-Kras* alleles in the absence and presence of Cre recombinase (CRE) to detect the unaltered wild-type *Kras* allele product (WT, 488 bp) as well as the unrecombined (*LSL-Kras*\*, 389 bp) and recombined (LoxP recombined, 616 bp) *LSL-Kras* allelic products. Gel images were cropped and color inverted for better visualization. Full-length gel images are provided in Figure 1***—source data 1***. **(C)** Levels of active KRAS determined by RBD pull-down (RBD-PD) followed by ELISA analysis using lysates derived from HEK-HT cells transiently expressing the indicated FLAG-tagged Kras proteins. Tubulin and empty vector serve as loading and negative controls, respectively. One of two biological replicates, see Figure 1—figure supplement 4C for the second biological replicate.

All four alleles were successfully recombined by Cre recombinase, as confirmed with two separate mouse embryonic fibroblast (MEF) cultures derived from each genotype (***Figure 1B***, *and* ***Figure 1—source data 1***). Based on common codons increasing protein expression and Q61R leading to higher levels of GTP-bound RAS, all four alleles displayed the expected gradual increase the level of active Kras. Namely, we confirm a stepwise increase in ectopic GTP-bound Kras in the ascending order of Kras^natG12D^<Kras^natQ61R^ <Kras^comG12D^<Kras^comQ61R^ by both immunocapture and ELISA-based assays (***Figure 1C*** and ***Figure 1—figure supplement 3***). Finally, we confirm that these four alleles exhibit the expected increase in oncogenic signaling *in vivo*. Namely, we crossed the *LSL-Kras^natG12D^/+*, *LSL-Kras^natQ61R^/+*, *LSL-Kras^comG12D^/+*, and *LSL-Kras^comQ61R^/+* genotypes into a *Rosa26-CreERT2* background, which expresses a tamoxifen-inducible Cre from the endogenous *Rosa26* promoter that is active in a broad spectrum of tissues (***Ventura et al., 2007***). Two adult mice from each of the four derived cohorts, as well as the control strain (*Rosa26-CreERT2/+*), were injected with tamoxifen and seven days later humanely euthanized, their lungs removed, RNA isolated, and the level of mRNA encoded by RAS target genes determined by qRT-PCR (***Figure 1—figure supplement 4A***). This revealed the expected increase in three of the target genes by the *LSL-Kras* alleles when encoded with common codons, which was further increased in the Q61R-mutant background (***Figure 1—figure supplement 4B,C***). Thus, the four novel *LSL-Kras* alleles function as designed.

### The effect of codon usage and mutation type on Kras biological activity

As the four alleles exhibited the expected biochemical and signaling properties of the encoded proteins, we next confirmed that these alleles were proportionally tumorigenic in a side-by-side comparison in the same organ. Each of these four alleles was specifically activated by tamoxifen injection to induce expression of Cre recombinase in the lung, after which every month thereafter for six months, five mice from each of the four cohorts were humanely euthanized (***Figure 2— figure supplement 1A***). The lungs from all mice were visually analyzed for the presence of surface pulmonary tumors (***Figure 2—figure supplement 1B***), and in addition, two H&E-stained sections from pairs of mice were assayed for the presence and type of pulmonary tumors by a veterinarian pathologist blinded to the genotype (***Figure 2A***). This revealed a stepwise increase in early onset and tumor burden of pulmonary lesions in lock step with the increased biochemical activity and signaling amplitude of the oncoproteins in the ascending order of Kras^natG12D^<Kras^natQ61R^ <Kras^comG12D^<Kras^comQ61R^ (***Figure 2A,B***). These tumorigenic phenotypes were the result of activating the *LSL-Kras* alleles in the lungs, as five control *Kras^+/+^* mice injected with tamoxifen failed to develop tumors after 13 months, twice the length of the study (***Figure 2—figure supplement 1C,D***). Finally, we find preliminary evidence that the G12D-mutant alleles preferentially led to atypical alveolar hyperplasia (AAH), whereas bronchiolar hyperplasia/dysplasia (BH) lesions were more prevalent with the Q61R-mutant alleles (***Figure 2— figure supplement 1E,F***). We conclude that this allelic set exhibits the expected increase in tumorigenic potential consistent with the activity and signaling of the encoded oncoproteins, but also exhibited evidence of mutation-specific effects on tumorigenesis.

**Figure 2.**
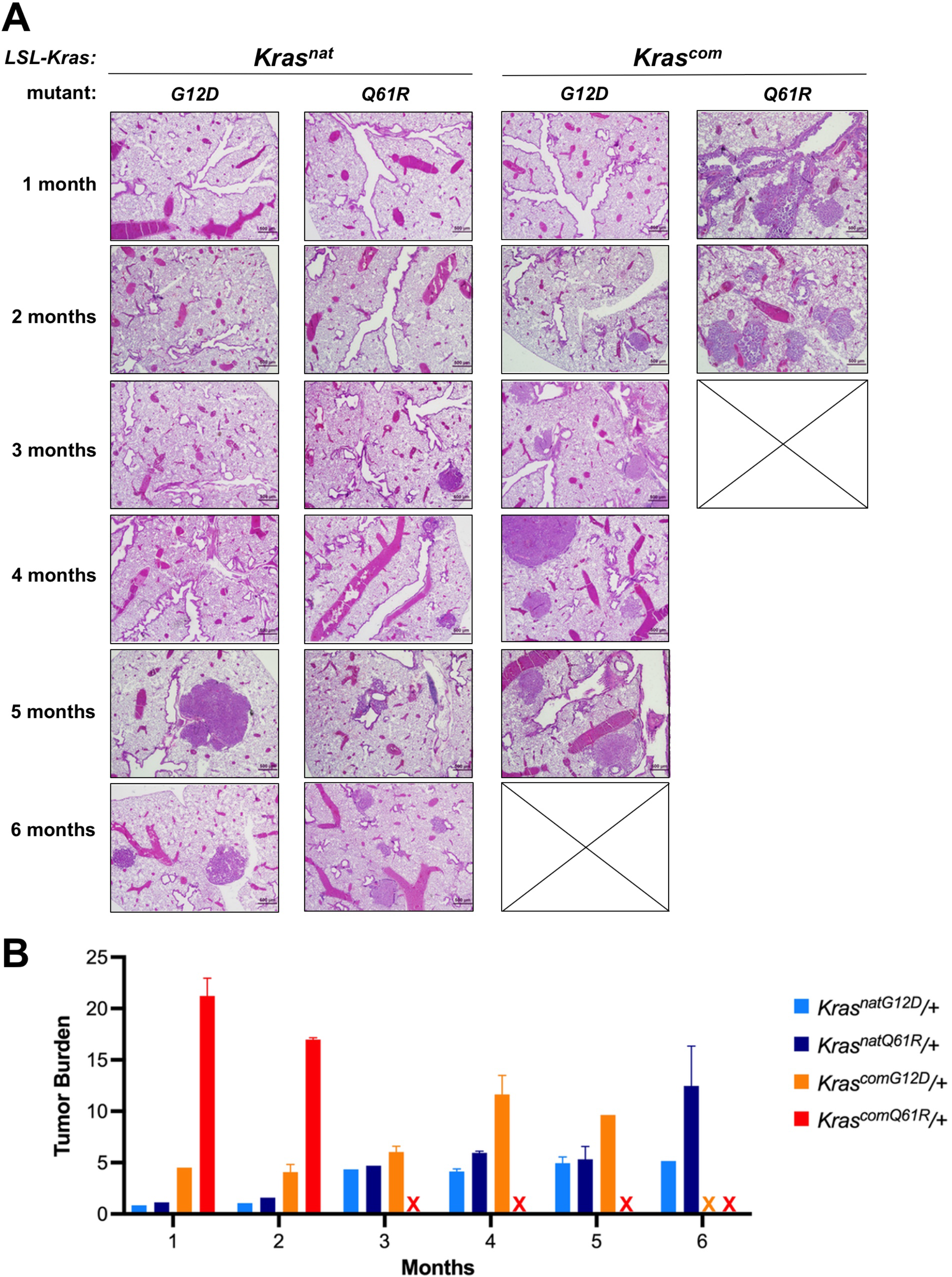
Biological effect of each oncogenic *LSL*-*Kras* allele. **(A)** Examples of H&E-stained lung sections, and **(B)** the mean *±* SD % tumor burden from microscopic analysis of two lung sections from five mice with the indicated *LSL-Kras* alleles in a *CC10-CreER/+* background at each of the indicated times post-tamoxifen injection.

### The response of normal cells to different Kras codon usage or mutations

As different tumor types and grades induced by each of these four oncogenic *LSL-Kras* alleles are presumably a product of tumor initiation, the key to understanding these effects must lie in how normal cells respond to these different oncoproteins. As the lung was sensitive to both the type of oncogenic mutation and the codon usage of the activated *Kras* allele, we compared the immediate transcriptional response to activation of each allele in this organ in the aforementioned *Rosa26-CreERT2/+* background. Cohorts of three adult male and female mice from each of the four cohorts were injected with tamoxifen, seven days later the animals were humanely euthanized, their lungs removed, and RNA isolated for bulk transcriptome sequencing (***Figure 3—figure supplement 1A*** and ***GSE181628***).

**Figure 3.**
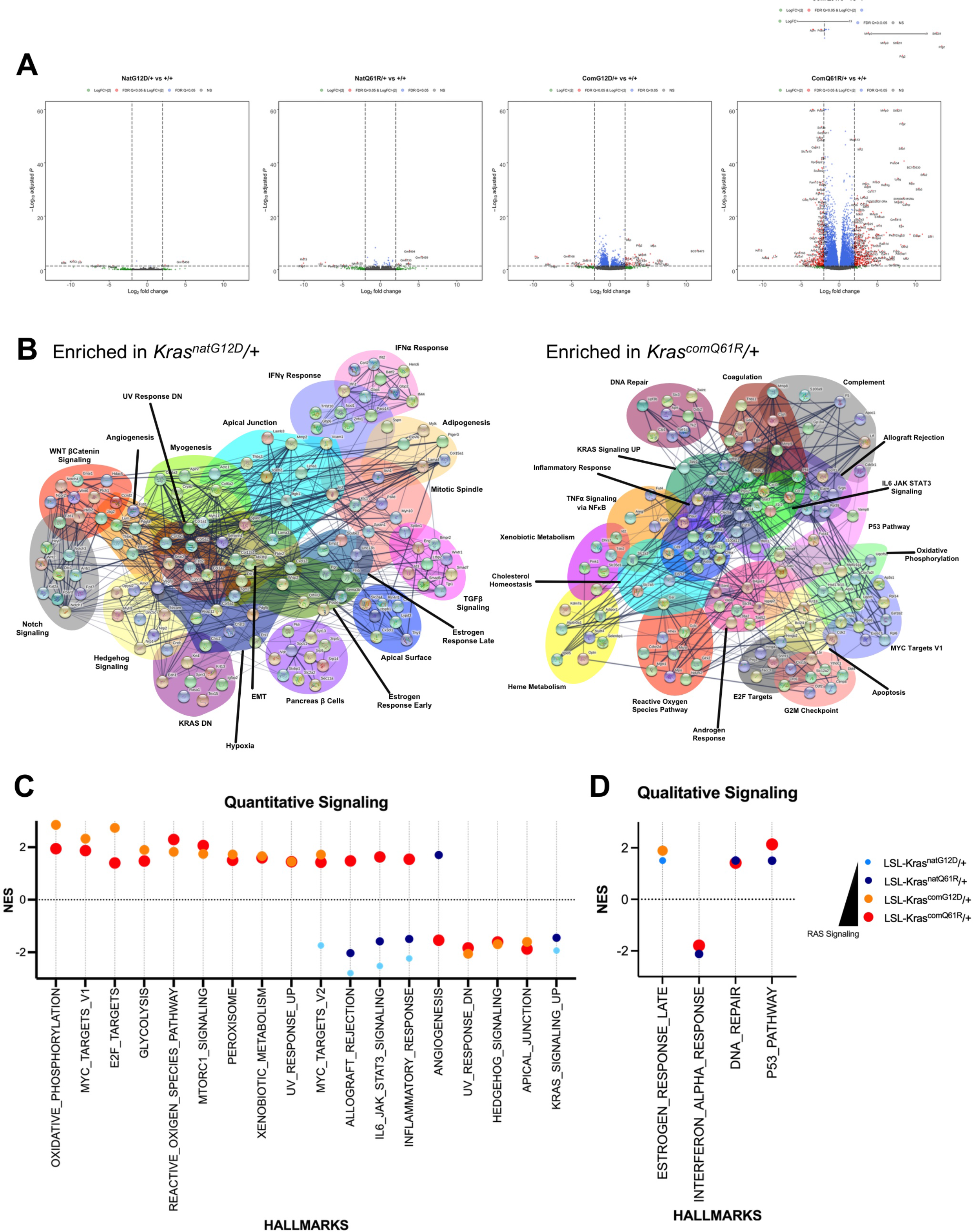
Transcriptomes of each oncogenic *LSL-Kras* allele. **(A)** Volcano plot of the log2 fold-change versus *p* value of the genes showing differential expression in each allele compared to the wild-type *Kras* allele in the lung. Full-sized plots are provided in Figure 3—figure supplement 2-5. **(B)** STRING analysis of the top ten genes in the differentially enriched GSEA hallmarks identified in RNA-seq analysis of the lungs of *Rosa26-CreERT2/+;LSL-Kras^natG12D^* versus *Rosa26- CreERT2/+;LSL-Kras^comQ61R^* mice seven days after tamoxifen injection. **(C,D)** Normalized enrichment score and patterns of the indicated GSEA hallmarks regulated by the quantitative **(C)** and qualitative **(D)** RAS signaling. Only hallmarks with an FDR less than 5% are shown. Dot size is adjusted to RAS activity for better visualization. All GSEA Hallmarks differentially regulated by quantitative and qualitative Ras signaling are provided in Figure 3— figure supplement 6.

This revealed that increasing Kras activity (through the combination of mutation type and codon usage) progressively moved the transcriptional signature away from one more similar to control tissue (*Kras^natG12D^*) to one much more complex and with features consistent with high oncogenic activity (*Kras^comQ61R^*) (***Figure 3A*** and ***Figure 3—figure supplement 2-5***). Focusing on the two extreme transcriptional responses, we repeated this analysis to ensure reproducibility, finding that activation of the *Kras^natG12D^* allele led to transcriptional signatures suggestive of the expansion of multipotent cells (***Figure 3—figure supplement 1B,C*** and ***GSE181627***). Namely, GSEA hallmarks indicative of Epithelial Mesenchymal Transition (EMT, TGF*β* Signaling, Wnt/*β*catenin Signaling, Notch Signaling, Apical Junction, Apical Surface, and Estrogen Response Early) and multiple cell lineages (Adipogenesis, Myogenesis, Hedgehog Signaling, and Pancreas *β* Cells). Conversely, activation of the *Kras^comQ61R^* allele had all features of a potent oncogenic signaling leading to oncogenic-induced stress. Namely, GSEA hallmarks indicative of high oncogenic signaling (KRAS Signaling UP) and unrestrained proliferation (MYC Targets V1, E2F Targets), leading to reactive oxygen species (Oxidative Phosphorylation, Reactive Oxygen Species Pathway) and a DNA damage response (DNA Repair, P53 Pathway, G2M Checkpoint) followed by apoptosis (Apoptosis) and inflammation (Inflammatory Response, IL6 JAK STAT3 Signaling, TNF*α* Signaling via NF-κB, Allograft Rejection). STRING analysis of the top ten genes from each of these GSEA hallmarks revealed clear crosstalk between the various signaling pathways, implying a consolidated signaling program by the two different oncoproteins manifests in very different responses by normal cells (***Figure 3B***). We thus suggest that this allelic set moves the response of normal cells from an expansion of multi-potent cells to one of extreme oncogenic signaling and stress.

To extract the effect of quantitative from qualitative signaling in this progression, we first performed principle component analysis on the four transcriptomes, which revealed that both *Kras^nat^* alleles were distinct from both *Kras^com^* alleles, while the G12D mutant was most distinct from the Q61R mutant in the *Kras^com^* background (***Figure 3—figure supplement 1D*** and ***GSE181628***). Next, we compared the transcriptional responses of both *Kras^nat^* alleles with both *Kras^com^* alleles to identify quantitative signaling differences, and compared both *Kras^G12D^* alleles with both *Kras^Q61R^* identify qualitative signaling differences. With regards to quantitative signaling, the two *Kras^com^* alleles induced signaling events related to oncogenic stress, such as an increase in Oxidative Phosphorylation, Glycolysis, Reactive Oxygen Species, MTORC1 signaling, Peroxisome, Xenobiotic Metabolism, and UV Response UP hallmarks and a decrease in UV Response Down, Hedgehog Signaling, and Apical Junction hallmarks. Conversely, the two *Kras^nat^* alleles decreased KRAS Signaling UP hallmarks (***Figure 3C*** and ***Figure 3—figure supplement 6***). We interpret this as expression level largely dictating the degree of oncogenic signaling in this allelic set. With regards to qualitative signaling, there were transcriptional signatures preferentially induced by specific mutations that were independent of codon usage. Namely, activation of both G12D-mutant alleles induced the Estrogen Response Late hallmark. On the other hand, we found that activation of the Q61R-mutant alleles induced DNA Repair and P53 Pathway and decreased Interferon Alpha Response hallmarks, suggesting that there was a strong tumor suppressive response uniquely activated by these mutant alleles (***Figure 3D*** and ***Figure 3—figure supplement 6***).

Finally, we validated the specific signaling responses detected by bulk RNAseq analysis by quantifying four to six marker genes in a subset of selected hallmarks. Namely, two adult mice from each of the four cohorts and control mice were injected with tamoxifen as above, and seven days later the animals were humanely euthanized, their lungs removed, RNA isolated, and the level of select transcripts determined by qRT-PCR. We identified similar expression patterns to the transcriptome signature in the hallmarks of TNF*α* Signaling via NFκB and Interferon-*γ*, which increase with Kras activity (***Figure 3—figure supplement 7A,B***), and EMT and Myogenesis, as these hallmarks were enriched in G12D mutants but depleted in Q61R mutants (***Figure 3—figure supplement 7A,C***).

Thus, in broad terms, we find that increasing Kras activation leads to progressively higher oncogenic signaling, eventually reaching the point of oncogenic stress, with the type of oncogenic mutation further modulating this response. Thus, quantitative signaling differences predominate between these mutants, with evidence of qualitative differences, potentially amplified at higher activation levels.

### Tissue sensitivities to different RAS signaling

With the biochemical, biological, and signaling output of each allele now determined, we next addressed the tropism of tissues towards specific oncogenic signaling by determining the tumor landscape upon globally activating each allele. As each of these alleles is expected to have different oncogenic potential, we opted for a moribundity endpoint, as opposed to a fixed endpoint, to identify the tissues most sensitive to tumorigenic conversion in a competition-based approach. To this end, each allele was again activated by tamoxifen using the aforementioned ubiquitous Cre driver *Rosa26-CreERT2*. Mice were then regularly monitored for moribundity endpoints, indicative of ensuing mortality due to cancer, at which point the animals were humanely euthanized (***Figure 4—figure supplement 1***). As a control to ruled out a tumor phenotype being a product of variations in gene activation, we validated Cre-mediated recombination between all four alleles across 14 diverse tissues seven days after tamoxifen injection, with only the ovary displaying reduced recombination, and hence was not included in the study (***Figure 4—figure supplement 2*, *Figure 4–source data 1***, and ***Supplementary File 1***).

**Figure 4.**
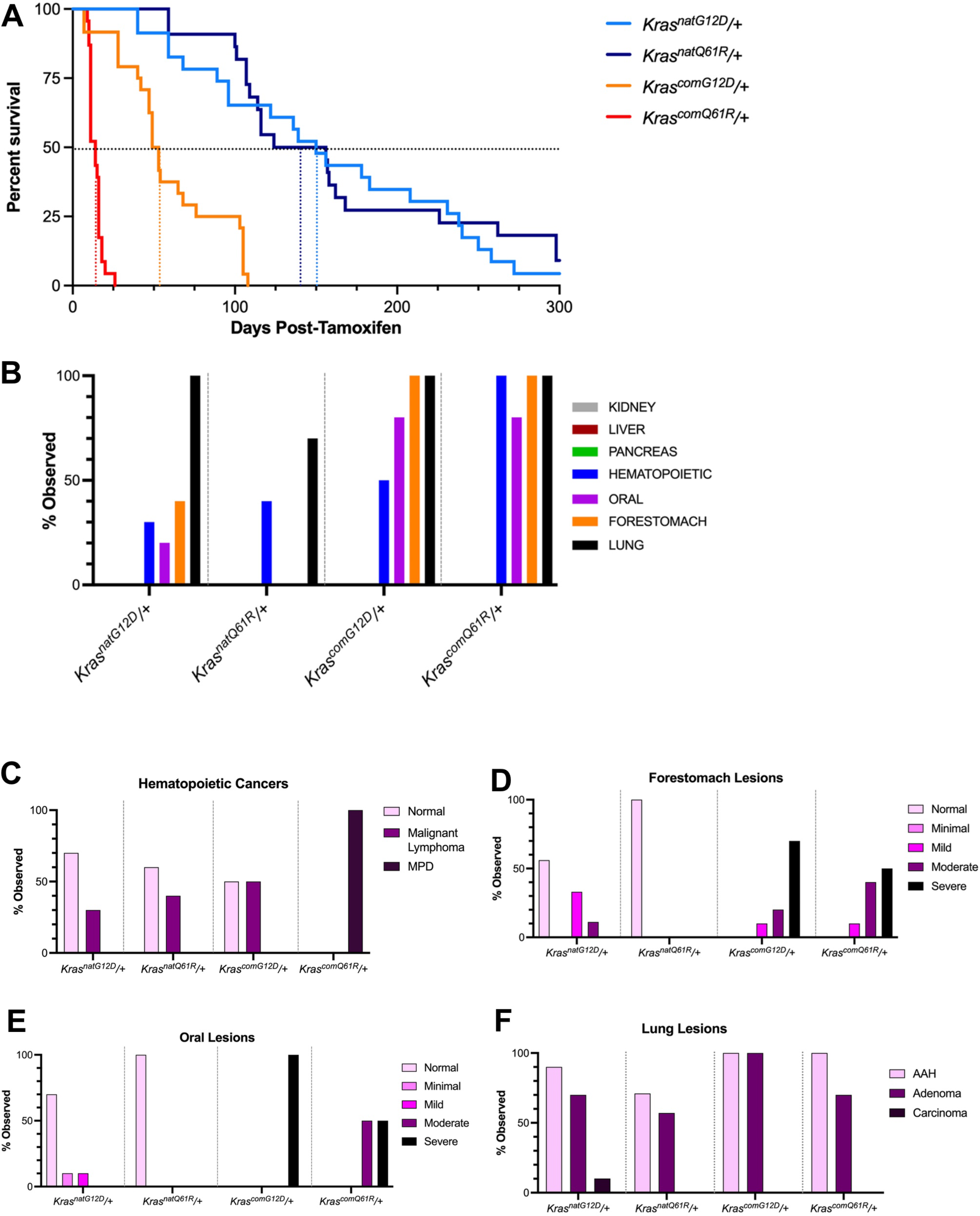
Tissue atlas of sensitivities to each oncogenic *LSL-Kras* allele. **(A)** Kaplan-Meier survival curve of the indicated *LSL-Kras* alleles after activation by tamoxifen. Dotted lines: 50% survival. **(B)** Number of mice with the indicated number of different tumor types. Examples of H&E-stained slides of the indicated tissues are provided in Figure 4—figure supplement 5. **(C-F)** Percentage of the indicated grades of **(C)** hematolymphopoietic, **(D)** forestomach, (**E**) oral, and **(F)** lung lesions at moribundity endpoint in *Rosa26-CreERT2/+* mice (*n*= 8 to 10) with one of the four indicated *LSL-Kras* alleles after activation by tamoxifen.

Plotting the percent survival for each of the four genotypes by the Kaplan-Meier approach revealed that the median lifespan of mice progressively increased from 14 days in the case of the activated *Kras^comQ61R^* allele to 150 days with the *Kras^natG12D^* allele (***Figure 4A***). Not surprisingly, the number of tumors per animal mirrored these survival differences (***Figure 4—figure supplement 3***). Pairwise comparisons of the survival curves revealed median survivals were statistically different, except between the *Kras^natG12D^* versus *Kras^natQ61R^* cohorts (***Figure 4—figure supplement 4*** and ***Supplementary file 2***). Of note, median survival was significantly decreased in mice with either of the *Kras^com^* alleles or with either of the Q61R mutants. As both common codons and the Q61R mutation increase Kras activity, tumorigenic potential, and signaling, we suggest that the level of active oncoprotein is the dominant determinant of cancer survival in this model.

To identify the tissue sensitivity to these different alleles, eight different organs were removed from the mice at necropsy and analyzed for pathologic changes as above. While there was much overlap, the prevalence and severity of specific cancer types varied between alleles, arguing that the different alleles can lead to differences in the tumor landscape (***Figure 4B*** and ***Figure 4—figure supplement 5***). This manifested in four general patterns of tissue sensitivity: hematolymphopoietic neoplasias increased in severity with the level of oncoprotein activation, squamous tumors were preferentially induced by the G12D-mutant alleles, carcinomas exhibited both these features, while many organs were resistant to oncogenic RAS-driven tumorigenesis within the timeframe of the study (***Figure 4A,B***, ***Figure 4—figure supplement 5***, and ***Supplementary file 3***). Collectively, these observations argue that all tested *LSL-Kras* alleles are oncogenic, with some tissues being sensitive and others resistant, and the response to these four alleles being variable, implying both qualitative and qualitative signaling differences contribute to the bias of tissue-specific oncogenicity of RAS mutations.

### A tissue sensitive to quantitative signaling

The incidence of hematolymphopoietic neoplasias increased with the biochemical activity, tumorigenic potential, and signaling amplitude of the oncoproteins. In more detail, the spleen and thymus from eight to ten mice from all four cohorts were removed at necropsy, after which H&E-stained sections were assayed for the presence and grade of hematolymphopoietic neoplasias as above. Beginning with the least active oncoprotein, most of the *Kras^natG12D^* mice had no evidence of hematolymphopoietic neoplasms, although some mice had pathologic features consistent with malignant lymphoma. The incidence of malignant lymphoma increased with *Kras^natQ61R^* allele, and then again with the *Kras^comG12D^* allele. Pathological analysis also revealed medullary hyperplasia in the thymus, as well as leukemic infiltrates in the kidneys and pancreas of these latter mice. Lastly, *Kras^comQ61R^* induced severe myeloproliferative disease (MPD) at 100% penetrance, with extensive myeloproliferative infiltrates throughout many tissues (***Figure 4B,C***, ***Figure 4—figure supplement 5***, ***Supplementary files 3,4***, and not shown). While we cannot discount high oncogenic activity leading to a different hematopoietic disease in these later mice, it seems more likely that this incredibly short latency for the onset of severe systemic myeloid neoplasia results in high mortality that prevents development of longer latency tumors such as the development of lymphopoietic neoplasms. Thus, with this proviso, hematolymphopoietic neoplasia are hyper-sensitive to quantitative oncogenic signaling, being induced at the lowest level of active Kras and progressively worsening with increased activity.

### Tissues sensitive to qualitative signaling

Proliferative lesions of forestomach and oral squamous epithelium exhibited evidence of qualitative signaling impacting tumorigenesis, namely tumors in these tissues were preferentially induced by the oncogenic G12D mutant of Kras encoded by either native or common codons. In the case of the forestomach tumors, pathological analysis performed as above revealed that the *Kras^natG12D^* allele induced squamous hyperplasia as well as mild and moderate grades of ‘atypical’ or dysplastic squamous lesions in the forestomach mucosa. Conversely, we did not detect any squamous proliferative changes in the *Kras^natQ61R^* mice (***Figure 4B,D***, ***Figure 4—figure supplement 5***, and ***Supplementary files 3,4***), despite the previously detected higher GTP-loading, tumorigenic potential, and signaling amplitude of *Kras^natQ61R^*. The same trend was observed in the *Kras^com^* background, except the difference was less extreme between the two mutants, and shifted towards more aggressive disease. Specifically, the *Kras^comG12D^* induced more severe grades of forestomach squamous lesions while *Kras^comQ61R^* now induced lesions that were of moderate grades (***Figure 4B,D***, ***Figure 4—figure supplement 5***, and ***Supplementary file 4***). Similar analysis of oral tumors revealed that activating the *Kras^natG12D^* allele induced minimal to mild grade squamous lesions, while the *Kras^natQ61R^* allele was not tumorigenic. Again, the *Kras^com^* background shifted disease to a more aggressive state. Namely, activating the *Kras^comG12D^* allele induced severe squamous papilloma in all mice, while the *Kras^comQ61R^* allele induced a mixture of moderate and severe grade squamous papillomas (***Figure 4B,E***, ***Figure 4—figure supplement 5***, and ***Supplementary file 4***). As such, in these two organs, the G12D mutant was associated with more severe phenotypes than the Q61R mutant, with changing codon usage to increase protein expression shifting this difference to a more advanced stage. These data collectively support a dominant role for qualitative differences in the tumorigenic sensitivity of the upper gastrointestinal epithelium, especially at lower activity levels.

### Tissues sensitive to both oncogenic activity and mutation type

The lung appeared to be a tissue whereby the effect of qualitative and quantitative signaling differences were both evident. In terms of qualitative signaling, pathological analysis performed as above revealed that the G12D mutation consistently induced more AAH and adenomas than the Q61R mutation in both the *Kras^com^* and *Kras^nat^* contexts (***Figure 4B,F***, ***Figure 4—figure supplement 5,6A***, and ***Supplementary file 4***). As was the case with forestomach and oral lesions, converting rare codons to common amplified the severity of lesions detected, which was particularly evident in the confluence of large peripheral AAH lesions induced preferentially by the G12D mutant allele (***Figure 4B,F***, ***Figure 4—figure supplement 5***, and ***Figure 4—figure supplement 6B***). In terms of quantitative signaling, no BH lesions were induced by either of the *Kras^nat^* alleles and instead were only prevalent in mice in which the *Kras^com^* alleles were activated. Further, the number of animals with BH lesions was higher upon activating the *Kras^comQ61R^* compared to the *Kras^comG12D^* allele (***Figure 4—figure supplement 5,6C***), akin to hematolymphopoietic neoplasia. Assuming that AAH and BH represent different types of tumors (and not different stages of the same tumor type), these data collectively support the lung being sensitive to both qualitative and quantitative oncogenic RAS signaling.

### Tissues resistant to oncogenic RAS driven tumorigenesis

Despite widespread tumorigenesis, we note that no overt proliferative lesions were detected at necropsy or by histopathologic analysis in the pancreas, kidney, or liver (***Figure 4—figure supplement 5*** and ***Supplementary file 3***). A gross survey of other organs such as the colon, intestine, heart, skin, and mammary glands also did not reveal any macroscopically detectable tumors (not shown). In agreement, many of these same tissues were reported to be refractory to tumorigenesis upon activation of an *LSL-Kras^G12D^* allele in the adult by CreER driven by a *Rosa26* (***Parikh et al., 2012***), *CK19* (***Ray et al., 2011***) or *Ubc9* (***Matkar et al., 2011***) promoter. Thus, many organs appear to be intrinsically resistant to the tumorigenic effects of oncogenic Kras, regardless of the mutation type or expression levels tested, at least within the timeframe of this study and in the tested *Rosa26-CreERT2* background.

## Discussion

Here we describe how separating oncogenic mutations from expression level in the endogenous *Kras* gene *in vivo* revealed quantitative and qualitative signaling contributions to RAS mutation patterns. We acknowledge three caveats to this approach. First, the four oncogenic *LSL-Kras* alleles were generated by fusing coding exons 1 to 3, an artificial gene architecture, and hence can only be compared to themselves and not to other types of *Kras* alleles. Second, these alleles were induced by an injection of tamoxifen to activate CreER expressed from the *Rosa26* locus, which is admittedly an unnatural situation whereby oncogenic Kras is expressed all at once in a multitude of cells and tissues, potentially perturbing homeostasis in the whole animal. Nevertheless, as the identical design was applied to all four alleles, comparisons can be made within this allelic set, and we find that targeted activation in the lung phenocopied activation by CreER expressed from the *Rosa26* locus. Third, Kras activity of this allelic set was defined in the lung. We therefore acknowledge that cell-type differences in feedback regulatory pathways (***Lake et al., 2016; Liu et al., 2018***), or codon-dependent expression (***Peterson et al., 2020***) could result in different levels of Kras expression, activation, or signaling compared to that observed in the lung.

With these above limitations in mind, transcriptome analysis shortly after activating each allele revealed a distinct cellular response. This allelic set began with the *Kras^natG12D^* allele, which induced EMT and differentiation hallmarks. EMT is known to promote stem cell-like fates (***Floor et al., 2011; Mani et al., 2008***) and activation of oncogenic Kras in the murine lung generates tumors with many tissue lineages (***Tata et al., 2018***). Taken together, we suggest that the transcriptional signature induced by the *Kras^natG12D^* oncoprotein reflects either reprogramming towards or an expansion of cells with multipotent characteristics. In support, single cell transcriptome profiling of lung tumors induced by targeted delivery of AAV-Cre in an *LSL- Kras^G12D^;Trp53^fl/fl^* background identified a distinct population with a mixed cellular identity (***Marjanovic et al., 2020***). At the other end of the spectrum, the *Kras^comQ61R^* allele induced transcriptional hallmarks consistent with overt oncogenic signaling. The transition from one transcriptional profile to a completely different profile appeared to be a product of both quantitative and qualitative signaling. As the level of Kras biochemical activity increased, so did the transcriptional signatures of RAS signaling, which at its crescendo, resulted in transcriptional signatures indicative of hyperproliferative and all the hallmarks of oncogenic stress. On the other hand, we also find evidence that G12D-mutant of Kras has a distinct signature compared to that of the Q61R mutant.

Globally activating these four alleles revealed four tumorigenic patterns. First, hematolymphopoietic neoplasias were largely driven by quantitative signaling, being induced at the lowest level of oncogenic activity and increasing in aggressiveness with increased Kras activity. To independently validate this result, we tested and found that globally activating a ‘super-rare’ version of *LSL-Kras^rareG12D^* allele encoded by the most rare codons possible induced hematolymphopoietic neoplasias, although in a much-protracted timeframe (***Figure 4—figure supplement 7*** and ***Supplementary file 3***), further supporting hematolymphopoietic tissues being hyper-sensitive to oncogenic Ras signaling. Second, proliferative lesions of forestomach and oral squamous epithelium were preferentially induced by G12D-mutant *Kras* alleles, pointing towards qualitative differences driving these tumors with higher expression shifting the effects to more aggressive grades. We acknowledge, however, that other interpretations beyond qualitative signaling differences between G12D and Q61R mutants are possible. It is worth mentioning that the survival is similar in mice in which *Kras^natG12D^* and *Kras^natQ61R^* alleles were globally activated, ruling out early moribundity masking the development of squamous lesion. Nevertheless, increasing oncoprotein expression does render the Q61R mutant tumorigenic in this tissue, so it is possible that qualitative signaling differences can be overcome by robust oncogenic activity. Third, the lung was sensitive to both forms of signaling, with qualitative (G12D-specific) signaling favoring AAH and adenomas while quantitative (Ras activity) favoring BH lesions, perhaps reflecting a different cell-of-origin for these different lesions (***Sutherland et al., 2014; Xu et al., 2014***). Fourth, with the two caveats that one, the short lifespan of some of the tested mice may prevent longer latency tumors from developing, and two, even though *Rosa26*-restricted Cre expression activated the four *Kras* alleles in tissues that did not form tumors, Cre may not be expressed in the tumor cell-of-origin within these tissues, we show that many organs appear to be intrinsically resistant to the tumorigenic effects of oncogenic Kras, at least within the timeframe of this study and in the *Rosa26-CreERT2* background. This suggests that the hematopoietic system, lungs, forestomach, and oral mucosa are unique in being sensitive to oncogenic Kras tumorigenesis, again however, within the confines of the experimental design. Interestingly, these tissue sensitivities share some similarity to that of humans, namely both species have RAS-associated cancers in the lung, mouth, and hematopoietic system but not in the mammary gland, skin, central nervous system and so forth, but there is also some discordance (***Prior et al., 2020***).

Our finding that oncogenic RAS mutations are not identical in terms of activity (GTP-loading), oncogenic potential (induction of lung tumorigenesis), and cellular response (transcriptome analysis), suggests the intriguing possibility that different initiating RAS mutations may have different therapeutic sensitivities. Indeed, censoring the GSEA hallmarks for pharmacological targets already drugged in the clinic identified FLT4/PDGFRB/KIT, EGFR, and ABL1/SRC as specific to *LSL-Kras^natG12D^* background, while AKT1/2, PI3KCA/CD/CG, SYK, RAF1, JAK2, TGFBR1, and CDK1/2/9 were specific to the *LSL-Kras^comQ61R^* background (***Figure 5***).

**Figure 5.**
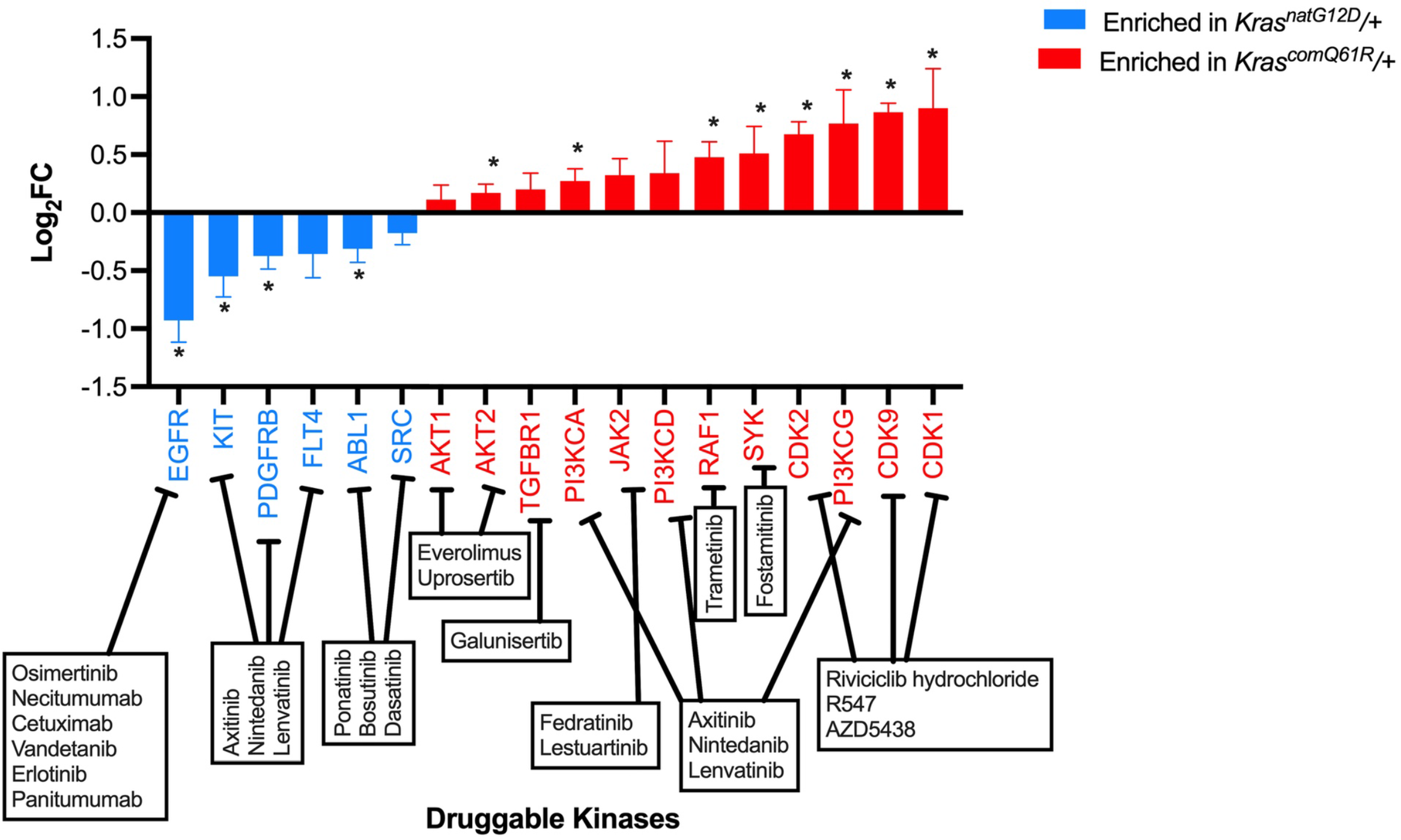
Transcriptome analysis predicts pharmacologic vulnerabilities of different oncogenic *LSL-Kras* alleles. Druggable kinases positively enriched in GSEA hallmarks identified in Figure 4B (blue *Kras^natG12D^*, red *Kras^comQ61R^*). * adjusted *p-*value less than 5%.

In summary, we interpret these results to imply that there are tissue sensitivities to quantitative and/or qualitative RAS signaling, with quantitative signaling favoring hematolymphopoietic neoplasias, qualitative signaling favoring squamous tumors, while carcinomas tended to be a combination of both types of signaling. The unique signaling dependencies of these tissues may, in turn, be capitalized upon to identify new therapeutic opportunities to target early tumorigenesis, when the tumors are particularly vulnerable, perhaps either as an early intervention or as a preventative measure in high-risk populations.

## Materials and methods

### Generation of LSL-Kras^natG12D^, LSL-Kras^natQ61R^, LSL-Kras^comG12D^, and LSL-Kras^comQ61R^ alleles

A bacteria artificial chromosome was engineered with 7.5 kbp of 5’ flanking sequence *Kras* intron 1 DNA, a Lox-STOP-Lox cassette (*LSL*) (***Feil et al., 1996***), exons 1 to 3 fused together and encoded by either native (*nat*) codons or with 95 rare codons converted to the most commonly used codons in the mouse genome (*com*) and either a G12D or Q61 oncogenic mutation, followed by the N-terminal 564-bp of intron 4, an FRT-Neomycin-FRT cassette, and a further 1.5 kbp of 3’ flanking sequence (***Figure 1A*** and ***Figure 1—figure supplement 1***). The targeting cDNAs were each cloned into the targeting PL253 vector (***Liu et al., 2003***) and electroporated into 129S6/C57BL/6N (G4) ES cells. After selection, clones were screened using PCR and positive clones were selected for each engineered *Kras* allele, expanded and frozen. At least two targeted clones per allele that are confirmed with a Southern hybridization were then microinjected into blastocysts to produce chimeras using standard procedures (***Behringer et al., 2014***). *LSL- Kras^natG12D^(+neo)*, *LSL-Kras^natQ61R^(+neo)*, *LSL-Kras^comG12D^(+neo)*, and *LSL-Kras^comQ61R^(+neo)* chimeras were crossed back to 129S6 mice. A genotyping PCR specific to the engineered *Kras* alleles with *neo* cassette was used to screen for germline transmission in clones. Each 129S6-*LSL- Kras^nat^(+neo)* and 129S6-*LSL-Kras^com^(+neo)* cohort was crossed with *ACTB:FLPe/+* (Jackson Laboratory, strain 003800) mice to remove the selection marker via FLP-mediated excision of the *neo* cassette (***Dymecki, 1996***). Removal of the *neo* cassette was confirmed with genotyping PCR. Resultant strains were backcrossed with 129S6 mice for five generations, generating the *LSL- Kras^natG12D^/+, LSL-Kras^natQ61R^/+*, *LSL-Kras^comG12D^/+*, and *LSL-Kras^comQ61R^/+* strains used in this study. All mouse care and experiments were performed in accordance with a protocol approved by the Institutional Animal Care and Use Committee (IACUC) of Duke University (Protocol no. A195-19- 09).

### Genotyping

For genotyping, genomic DNA was isolated from 1-2 mm piece of toes by boiling for 30 min in 100 *μ*l Toe Lysis Buffer (25 mM NaOH and 0.2 mM EDTA), followed by neutralization with 1.5 volume of Neutralization Buffer (40 mM Tris-HCl pH 6.0). 1.2 *μ*l of the isolated genomic DNA was subjected to PCR. Genotyping cell lines and mouse tissue was performed as above with 20 ng of genomic DNA isolated with a Qiagen DNeasy Blood and Tissue DNA Extraction Kit (Qiagen, #69504). All genotyping PCR reactions were performed using 0.4 U (0.08 *μ*l) Platinum Taq polymerase (Invitrogen, #10342046) in 12.5 *μ*l reaction volume with final concentration of 1.5 - 2 mM MgCl2, 0.2 mM of each dNTP, and 0.5 *μ*M of each primer. Full-length gels and replicates are provided (***Figure 1—source data 1*** and ***Figure 4—source data 1***). Mice were genotyped using the following primers:

*LSL-Kras(+neo)* alleles:

Kras.in3.F: *5*’-TTGGTGTACATCACTAGGCTTCA-*3*’ Kras.in3.R: *5*’-TGGAAAGAGTAAAGTGTGGTGGT-*3*’ Kras.neo.F: *5*’-GTGGGCTCTATGGCTTCTGA-*3*’

Products: 590 bp (Targeted Allele[+neo]) or 240 bp (WT allele)

*LSL-Kras^com^* alleles:

KrasCOM5.F: *5*’-CTTCCATTTGTCACGTCCTGC-*3*’ KrasCOM5.R: *5*’-TCTTCGGTGGAAACAACGGT-*3*’

Product: 448 bp (LSL-Krascom)

*LSL-Kras^nat^* alleles:

F-LSL: *5*’-TAGTCTGTGGGACCCCTTTG-*3*’ R-LSL: *5*’-GCCTGAAGAACGAGATCAGC-*3*’

Product: 448 bp (LSL-Krasnat) Recombination PCR:

KRASOP.A2: *5*’-CTAGCCACCATGGCTTGAGT-*3*’ KRASOP.B: *5*’-GTAATCACAACAAAGAGAATGCAG-*3*’ LSL-F: *5*’-GGGGAACCTTTCAGGCTTA-*3*’

Products: 616 bp (LoxP Recombined), 488 bp (WT allele), or 389 bp (*LSL* allele)

*CC10-CreER* alleles:

F-CC10 WT: *5*’-ACTCACTATTGGGGGTGTGG-*3*’ R-CC10 WT: *5*’-GGAGGACTTGTGGATCTTG-*3*’ F-Cre: *5*’-TCGATGCAACGAGTGATGAG-*3*’

R-Cre: *5*’-TTCGGCTATAGGTAACAGGG-*3*’

Products: 450 bp (*CC10-CreER* allele) or 350 bp (WT allele)

*Rosa26-fGFP* alleles:

F-Rosa-01: 5’-CACTTGCTCTTCCAAAGTCG-*3*’ R-Rosa-02B: *5*’-TAGTCTAACTCGCGACACTG-*3*’ F-CAG-02B: *5*’-GTTATGTAACGCGGAACTCC-*3*’

Products: 500 bp (WT allele allele) or 350 bp (*Rosa26-fGFP* allele)

*Rosa26-CreERT2* alleles:

R26R-univF: *5*’-AAAGTCGCTCTGAGTTGTTAT-*3*’ R26R-wtR: *5*’-GGAGCGGGAGAAATGGATATG-*3*’ CreER-R1: *5*’-CCTGATCCTGGCAATTTCG-*3*’

Products: 800 bp (*Rosa26-CreERT2* allele) or 600 bp (WT allele)

*LSL-Kras* (LSL element):

KRASOP.A2: *5*’-CTAGCCACCATGGCTTGAGT-*3*’ KRASOP.B: *5*’-GTAATCACAACAAAGAGAATGCAG-*3*’

Product: 389 bp (*LSL-Kras* alleles) CreER validation:

F-Cre: 5’-GGAGGACTTGTGGATCTTG-*3*’ CREER-R1: *5*’-CCTGATCCTGGCAATTTCG-*3*’

Product: 500 bp (CreER)

*LSL-Kras^rare^* alleles:

KrasRAR.F: *5*’-TATGCGTACGGGTGAAGGTT-*3*’ KrasRAR.R: *5*’-GCAGAGCACAGACTCACGTC-*3*’

Product: 275 bp (*LSL-Kras^rare^*)

### Plasmids

N-terminal FLAG-tagged *Kras* cDNAs *FLAG-Kras^natG12D^, FLAG-Kras^natQ61R^, FLAG-Kras^comG12D^,* and *FLAG-Kras^comQ61R^* were designed using the same sequence of exons 1-3 as described in the engineered alleles with the addition of the native *Kras4B* to the C-terminus and cloned into pcDNA3.1+ (Thermo-Fisher). The plasmid sequences were verified by sequencing.

### Mouse embryonic fibroblasts (MEFs)

*LSL-Kras^natG12D^/+*, *LSL-Kras^natQ61R^/+, LSL-Kras^comG12D^/+*, and *LSL-Kras^comQ61R^/+* mice were crossed with 129S6 mice for timed pregnancies to isolate embryos 12.5-13.5 days postcoitum, as previously described (***Serrano et al., 1997***). Two primary MEF clones from each cohort cultured in DMEM with high glucose (Sigma-Aldrich, D5796) supplemented with 10% FBS (VWR, 97068-085) were immortalized by infection with an amphotropic retrovirus derived from plasmid pBABE-neo largeTcDNA (a gift from Robert Weinberg, Addgene plasmid #1780) (***Hahn et al., 2002***) using standard methodologies (***O’Hayer & Counter, 2006***). Paired parental MEF lines that were not infected with the pBABE-neo largeTcDNA was used as a negative control for antibiotic selection. Stable immortalized MEF cultures were selected in 400 μg/ml Geneticin (Thermo- Fisher, 10131035) and again infected with an amphotrophic retrovirus derived from plasmid MSCV-Cre-Hygro (a gift from Kai Ge, Addgene plasmid #34565) to activate the mutant *Kras* allele (***Wang et al., 2010***). The stable Cre-expressing MEF cultures were selected in 500 μg/ml hygromycin (Thermo-Fisher, 10687010). Paired parental MEF lines that were not infected with MSCV-Cre-Hygro were used as negative control for hygromycin selection and for recombination PCR. Genomic DNA was isolated from the resultant cells using Qiagen DNeasy Blood and Tissue DNA Extraction Kit (Qiagen, #69504) and subjected to PCR to detect recombination of the *LSL-Kras* alleles as described above. Full-length gel image is provided (***Figure 1—source data 1***).

### Ectopic expression, immunoblots, and Ras activity assay

To validate the expression levels of FLAG-tagged *Kras* constructs 2 x 10^6^ HEK-HT cells (***Counter et al., 1992***) were seeded in 10 cm tissue culture plates in DMEM with high glucose (Sigma-Aldrich, D5796) supplemented with 10% FBS (VWR, 97068-085), and were transiently transfected the next day with the pcDNA3.1+ (empty vector) or the same plasmid encoding *FLAG-Kras^natG12D^*, *FLAG-Kras^natQ61R^*, *FLAG-Kras^comG12D^*, or *FLAG-Kras^comQ61R^* using FuGene 6 reagent (Promega, E2691) according to the manufacturer’s protocol. 48 h later, transfected cells were washed with cold PBS and pelleted. Cell pellets were lysed in 5 volumes of 1x lysis buffer (50 mM Tris-HCl pH 8.0, 150 mM NaCl, 1% NP40, 0.5% sodium deoxycholate, 0.1% SDS, and 5 mM EDTA) containing Protease Inhibitor Cocktail (Roche, 11836170001). The DC Protein Assay (Bio-Rad, 5000112) was used to measure protein concentration and 30 *μ*g protein of each sample was resolved by SDS-PAGE, transferred to a PVDF membrane (Bio-Rad, 1704273), blocked in 5% milk, and immunoblotted by the following antibodies in 5% BSA (Sigma, A7906-500G): FLAG (Sigma, F1804; diluted 1:1000), KRAS (Santa Cruz, SC-30; diluted 1:500), and ß-Tubulin (Sigma, T5201; diluted 1:10000). Primary antibody incubation was performed at room temperature for 1 h followed by the secondary antibody incubation at 4*°*C overnight. To measure biochemical activity of the FLAG-tagged *Kras* constructs, cells were seeded, transfected, washed with PBS as above, after which cell pellets were lysed and immediately assayed with the Active Ras Detection Kit (Cell Signaling, #8821) or Ras GTPase ELISA Kit (Abcam, ab134640) according to the manufacturer’s protocol. Measurements were done with two technical replicates of four serial dilutions and data shown are representative of two independent biological replicates. Full-length images of immunoblots and replicates are provided (***Figure 1—source data 2***).

### Tumorigenesis studies

LSL-Kras^natG12D^/+, LSL-Kras^natQ61R^/+, LSL-Kras^comG12D^/+, and LSL-Kras^comQ61R^/+ mice were crossed with either CC10-CreER/+;R26-CAG-fGFP/+ (a gift from Mark Onaitis) (**Xu et al., 2012**) or Rosa26- CreERT2/+ (Jackson Laboratory, strain 008463) mice and animals with the desired alleles were selected by genotyping. At 6 to 8 weeks of age selected littermates with random distribution of males and females received four intraperitoneal injections of tamoxifen (Sigma-Aldrich, T5648- 5G, CAS# 10540-29-1) dissolved in corn oil (Sigma-Aldrich, C8267) and filter sterilized for a dose of 250 μg/g body weight to induce Cre-mediated recombination of the LSL cassette, activating the engineered oncogenic mutant Kras alleles. Mice from the first cross were humanely euthanized 1, 2, 3, 4, 5, or 6 months later while mice from the second cross were humanely euthanized upon moribundity. To prevent age-related outcomes, four mice that didn’t reach moribundity by 300 days were humanely euthanized at one year of age, which we reasoned to suffice to show tumorigenesis as it is about 8 times longer than the previously identified average moribundity of 5-6 weeks with the tamoxifen treated Rosa26-CreERT2/+; LSL-KrasG12D/+ mice (**Parikh et al., 2012**). Selected tissues were removed at necropsy and fixed in 10% Formalin (VWR, 89370-094) for 24 to 48 h, then post-fixed in 70% ethanol (VWR, 89125-166) until analysis.

### Tissue analysis

Animals were humanely euthanized with inhaled carbon dioxide and subjected to a complete necropsy. Selected organs were sampled for microscopic examination including lung, liver, kidney, spleen, thymus, stomach, pancreas, and macroscopic lesions. All tissues were fixed for 48 h in 10% neutral buffered formalin (VWR, 89370-094) and then post-fixed in 70% ethanol (VWR, 89125-166), processed routinely, embedded in paraffin with the flat sides down, sectioned at a depth of 5 *μ*m, and stained by the H&E method. Routine processing of the lungs from *CC10- CreER/+*;*R26-CAG-fGFP/+* mice was performed by the Duke Research Immunohistology Lab, while all tissues from *Rosa26-CreERT2/+* mice were processed by IDEXX Laboratories. Tissues and H&E slides were evaluated by a board certified veterinary anatomic pathologist with experience in murine pathology.

### Tissue recombination analysis

At 6 to 8 weeks of age, two female mice from each of the genotypes *LSL-Kras^natG12D^/+, LSL- Kras^natQ61R^/+, LSL-Kras^comG12D^/+,* and *LSL-Kras^comQ61R^/+* in a *Rosa26-CreERT2/+* background were injected with tamoxifen as above, and seven days later humanely euthanized. One age-matched female *Rosa26-CreERT2/+; LSL-Kras^comG12D^* mouse was also humanely euthanized as a no-tamoxifen control. At necropsy the colon, duodenum, ileum, cecum, jejunum, pancreas, spleen, glandular stomach, forestomach, kidney, liver, lung, heart, and ovaries were removed, genomic DNA extracted, and the status of recombination of each of the four oncogenic mutant *LSL-Kras* alleles determined by recombination PCR in duplicate, as described above. The intensities of bands corresponding to the unaltered wild-type *Kras* allele (WT) as well as the unrecombined and recombined oncogenic mutant *LSL-Kras* alleles was quantified with ImageJ software and the recombination rates for each tissue type from each mouse calculated by dividing the densitometry of recombined allele to that of the *WT Kras* allele. Full-length gels and replicates are provided (***Figure 4—source data 1***).

### Codon usage plots

The codon usage index (***Sharp & Li, 1987***) was calculated using the relative codon frequency derived from codon usage in the mouse exome (***Nakamura et al., 2000***) with a sliding windows of 25 codons across the open reading frame (ORF) of each transcript. A theoretical *Kras* ORF encoded by the rarest codons at each position (grey dotted line) was plotted for reference (***Figure 1—figure supplement 1***).

### Statistical analysis

Statistical analyses were performed using GraphPad Prism software, version 8 (GraphPad Software). One-way ANOVA with Bonferroni’s multiple comparisons test with a single pooled variance and a 95% CI were used for experiments with more than 2 groups. Reported *P* values are adjusted to account for multiple comparisons. A *P* value of less than 0.05 was considered statistically significant.

### RNAseq

Three mice with random distribution of males and females were selected from cohorts of *LSL- Kras^natG12D^/+, LSL-Kras^natQ61R^/+, LSL-Kras^comG12D^/+,* and *LSL-Kras^comQ61R^/+* mice in a *Rosa26- CreERT2/+* background, injected with tamoxifen at 6 to 8 weeks of age as described above, and humanely euthanized seven days later to harvest lungs. Three mice with random distribution of males and females in *Rosa26-CreERT2/+* background without the *LSL-Kras* alleles were used as negative control. For the second RNAseq experiment, three mice with random distribution of males and females were selected from *LSL-Kras^natG12D^/+, and LSL-Kras^comQ61R^/+* mice in a *Rosa26- CreERT2/+* background, injected with tamoxifen at 6 to 8 weeks of age as described above, and humanely euthanized seven days later to harvest lungs. All tissue lysis and RNA extraction steps were performed in a chemical hood and all instruments and tools were sprayed with RNaseZAP™ (Sigma, R2020) to prevent RNA degradation. Isolated lungs were immediately stored in RNA stabilizing solution RNAlater (Sigma, R0901) at 4*°*C overnight and then transferred to -80*°*C for long-term storage. Lung tissues were thawed, weighed, and pulverized with mortar and pestle in the presence of liquid nitrogen. RNA extraction was performed immediately thereafter using RNAeasy Kit (Qiagen, 74104) according to manufacturer’s instructions. Briefly, tissue lysates were prepared according to the kit instructions and tissue clumps were removed using QIAshredder columns (Qiagen, 79654). Cleared lysates were applied to RNeasy silica columns and on-column DNase digestion was performed using RNase-Free DNase Set (QIAGEN, 79254) to remove DNA in silica membrane. Following wash steps, RNA was eluted in RNase-free water that was treated with RNAsecure™ RNase Inactivation Reagent (Thermo-Fisher, AM7005) according to manufacturer’s instructions. Extracted total RNA quality and concentration was assessed on Fragment Analyzer (Agilent Technologies) and Qubit 2.0 (ThermoFisher Scientific), respectively. Samples with RIN less than 7 were not sequenced. RNA-seq libraries were prepared using the commercially available KAPA Stranded mRNA-Seq Kit (Roche). In brief, mRNA transcripts were captured from 500 ng of total RNA using magnetic oligo-dT bead. The mRNA was then fragmented using heat and magnesium, and reverse transcribed using random priming. During second strand synthesis, the cDNA:RNA hybrid was converted into to double-stranded cDNA (dscDNA) and dUTP was incorporated into the second cDNA strand, effectively marking this strand. Illumina sequencing adapters were ligated to the dscDNA fragments and amplified to produce the final RNA-seq library. The strand marked with dUTP was not amplified, allowing strand-specificity sequencing. Libraries were indexed using a dual indexing approach allowing for all the libraries to be pooled and sequenced on the same sequencing run. Before pooling and sequencing, fragment length distribution for each library was first assessed on a Fragment Analyzer (Agilent Technologies). Libraries were also quantified using Qubit. Molarity of each library was calculated based on qubit concentration and average library size. All libraries were then pooled in equimolar ratio and sequenced. Sequencing was done on an Illumina NovaSeq 6000 sequencer. The pooled libraries were sequenced on one lane of an S-Prime flow cell at 50bp paired-end. Once generated, sequence data was demultiplexed and Fastq files generated using bcl2fastq v2.20.0.422 file converter from Illumina. The RNAseq data has been deposited in NCBI’s Gene Expression Omnibus (***Edgar et al., 2002***). Initial and secondary RNAseq data are available through GEO Series accession numbers ***GSE181628*** and ***GSE181627***, respectively, and are accessible under the project ***GSE181629*** (https://www.ncbi.nlm.nih.gov/geo/query/ acc.cgi?acc=GSE181629).

### Transcriptome analysis

RNA-seq data was processed using the TrimGalore toolkit (http://www.bioinformatics. babraham.ac.uk/projects/trim_galore) which employs Cutadapt (***Martin, 2011***) to trim low-quality bases and Illumina sequencing adapters from the 3’ end of the reads. Only reads that were 20 nucleotides or longer after trimming were kept for further analysis. Reads were mapped to the GRCm38v73 version of the mouse genome and transcriptome (***Kersey et al., 2012***) using the STAR RNA-seq alignment tool (***Dobin et al., 2013***). Reads were kept for subsequent analysis if they mapped to a single genomic location. Gene counts were compiled using the HTSeq tool (http://www-huber.embl.de/users/ anders/HTSeq/). Only genes that had at least 10 reads in any given library were used in subsequent analysis. Normalization and differential expression was carried out using the DESeq2 (***Love et al., 2014***) Bioconductor (***Huber et al., 2015***) package with the R statistical programming environment (http://www.R-project.org). The false discovery rate was calculated to control for multiple hypothesis testing (***Figure 3—figure supplement 2-5***). Gene set enrichment analysis (***Mootha et al., 2003***) was performed to identify hallmarks and pathways associated with altered gene expression for each of the comparisons performed (***Figure 3—figure supplement 6*** and ***Figure 3—source data 1***). To identify low Kras- and high Kras-specific druggable kinases, the transcriptome enriched in *Kras^natG12D^/+ and Kras^comQ61R^/+*- specific GSEA hallmarks were cross referenced with kinases whose clinical inhibitors were previously surveyed (***Klaeger et al., 2017***). The kinases enriched in *Kras^natG12D^/+* and in *Kras^comQ61R^/+* are shown in blue and red, respectively with adjusted *p-*value less than 5% were highlighted (***Figure 5***).

### Quantitative Real-Time PCR

Two mice with random distribution of males and females were selected from cohorts of *LSL- Kras^natG12D^/+, LSL-Kras^natQ61R^/+, LSL-Kras^comG12D^/+,* and *LSL-Kras^comQ61R^/+* mice in a *Rosa26- CreERT2/+* background, injected with tamoxifen at 6 to 8 weeks of age as described above, and humanely euthanized seven days later to harvest lungs. Three mice with random distribution of males and females in *Rosa26-CreERT2/+* background without the *LSL-Kras* alleles were used as negative control. RNA extraction was performed as mentioned above, followed by first strand cDNA synthesis from 2 *μ*g RNA, and real-time PCR using GoTaq 2-Step RT-qPCR kit (Promega, A6110). All measurements were normalized against Actin as the internal control using the 2^-^*^ΔΔ^*^Ct^ method (***Figure 1—source data 3*** and ***Figure 3—source data 2***). For qRT-PCR analysis for the second biological replicate, one of two negative controls did not have numerical data for two transcripts and hence was not used for plotting. Data shown are representative of two independent biological replicates and the primer sequences are:

GLI1-F: 5’-CCCATAGGGTCTCGGGGTCTCAAAC-3’, GLI1-R: 5’-GGAGGACCTGCGGCTGACTGTGTAA-3’ CDH1-F: 5’-GTCTCCTCATGGCTTTGC-3’, ECAD-R: 5’-CTTTAGATGCCGCTTCAC-3’

TWIST1-F: 5’-AGCGGGTCATGGCTAACG-3’, TWIST1-R: 5’-GGACCTGGTACAGGAAGTCGA-3’ ZEB2-F: 5’-GAGCTTGACCACCGACTC-3’, ZEB2-R: 5’-TTGCAGGACTGCCTTGAT-3’

SOX5-F: 5’-ATTGTGCAGTCCCACAGGTTG-3’, SOX5-R: 5’-CTGCCTTTAGTGGGCCAGTG-3’ IL6-F: 5’-CCGGAGAGGAGACTTCACAG-3’, IL6-R: 5’-CAGAATTGCCATTGCACAAC-3’ IL10-F: 5’-GGTTGCCAAGCCTTATCGGA-3’, IL10-R: 5’-ACCTGCTCCACTGCCTTGCT-3’ TNFA-F 5’-CCCCAAAGGGATGAGAAGTT-3’, TNFA-R: 5’-GTGGGTGAGGAGCACGTAGT-3’ CCL2-F: 5’-AGGTCCCTGTCATGCTTCTG-3’, CCL2-R: 5’-TCTGGACCCATTCCTTCTTG-3’ LIF-F: 5’-AATGCCACCTGTGCCATACG-3’, LIF-R: 5’-CAACTTGGTCTTCTCTGTCCCG-3’

TNFSF9-F: 5’-GCAAGCAAAGCCTCAGGTAG-3’, TNFSF9-R: 5’-TCCAGGAACGGTCCACTAAC-3’ IFNG-F: 5’-CGGGAGGTGCTGCTGATGG-3’, IFNG-R: 5’-AGGGACAGCCTGTTACTACC-3’

### Generation of *LSL-Kras^rareG12D^/+* mice and tumorigenesis study

An additional inducible *Kras* allele with the most rare codons was generated in the same manner as mentioned above for the four *LSL-Kras* alleles from BAC design to chimera production (***Figure 1—figure supplement 1***). *LSL-Kras^rareG12D^(+neo)* chimeras were crossed back to 129S6 mice followed by *ACTB:FLPe/+* (Jackson Laboratory, strain 003800) mice to remove the *neo* selection marker via FLP-mediated excision (***Dymecki, 1996***). Both germline transmission and the removal of the *neo* cassette were confirmed with genotyping PCR as mentioned above. Resultant strain was backcrossed with 129S6 mice for five generations, generating the *LSL-Kras^rareG12D^/+* strain. *LSL-Kras^rareG12D^/+* mice were crossed with *Rosa26-CreERT2/+* (Jackson Laboratory, strain 008463) mice and animals with the desired alleles selected by genotyping. At 6 to 8 weeks of age seven male and female adult mice received four intraperitoneal injections of tamoxifen (Sigma-Aldrich, T5648-5G, CAS# 10540-29-1) dissolved in corn oil (Sigma-Aldrich, C8267) and filter sterilized for a dose of 250 *μ*g/g body weight to activate the engineered oncogenic mutant *Kras* allele. Tamoxifen injected mice were humanely euthanized at the fixed timepoint of 22 months later with inhaled carbon dioxide. Lung, liver, kidney, spleen, thymus, stomach, pancreas, and femurs were removed at necropsy and fixed in 10% Formalin (VWR, 89370-094) for 24 to 48 h, then post-fixed in 70% ethanol (VWR, 89125-166), processed routinely, embedded in paraffin with the flat sides down, sectioned at 5 *μ*m, and stained with H&E. Routine processing of the tissues was performed by IDEXX Laboratories. Tissues and H&E slides were evaluated by a board certified veterinary anatomic pathologist with experience in murine pathology.

## Online Supplemental Material

***Figure 1—figure supplement 1*** shows the codon usage index versus codon number of *Kras* with native, common, and rare codons. ***Figure 1—figure supplement 2*** shows the sequence alignment of the coding exons of the four novel *LSL-Kras* alleles used in this study. ***Figure 1—figure supplement 3*** shows protein expression and activity levels of the encoded *Kras* proteins. ***Figure 1—figure supplement 4*** shows the qRT-PCR analysis of the RAS target genes. ***Figure 2—figure supplement 1*** shows the lung-specific activation of the four *LSL-Kras* alleles. ***Figure 3—figure supplement 1*** shows transcriptomic analysis of the lung immediately after activation of the four *LSL-Kras* alleles. ***Figure 3—figure supplement 2-5*** shows the volcano plot of the *log2* fold-change versus *p* value of differentially expressed genes from RNA-seq analysis of the lungs of *Rosa26- CreERT2/+;Kras+/+* mice versus ***figure supplement 2*** *Rosa26-CreERT2/+;LSL-Kras^natG12D^/+, **figure supplement 3** Rosa26-CreERT2/+;LSL-Kras^natQ61R^/+ **figure supplement 4** Rosa26-CreERT2/+;LSL- Kras^comG12D^/+,* and ***figure supplement 5*** *Rosa26-CreERT2/+;LSL-Kras^comQ61R^/+* mice. ***Figure 3— figure supplement 6*** shows GSEA hallmarks regulated by the four *LSL-Kras* alleles. ***Figure 3— figure supplement 7*** shows validation of the transcriptome via qRT-PCR analysis of selected genes. ***Figure 4—figure supplement 1*** shows the experimental design to determine tissue sensitivities of the four *LSL-Kras* alleles. ***Figure 4—figure supplement 2*** shows the recombination analysis of 14 organs after tamoxifen injection. ***Figure 4—figure supplement 3*** shows the number of different tumor types detected in each allele at the moribundity endpoint. ***Figure 4— figure supplement 4*** shows pairwise comparisons of the survival curves based on Log-Rank (Mantel Cox) test. ***Figure 4—figure supplement 5*** shows the examples of H&E-stained tissues from multiple organs from each of the four *LSL-Kras* alleles. ***Figure 4—figure supplement 6*** analysis of pulmonary lesions in the *Rosa26-CreERT2/+* background. ***Figure 4—figure supplement 7*** shows the tumor landscape after activating the ‘super-rare’ *LSL-Kras^rareG12D^* 20 months after tamoxifen injection.

***Figure 1—source data 1*** shows the full-length gel images of genotyping of MEF cultures in ***Figure 1B***. ***Figure 1—source data 2*** shows the full-length gel images of immunoblots in ***Figure 1—figure supplement 3***. ***Figure 1—source data 3*** shows shows the Ct values from the qRT-PCR analysis in ***Figure 1—figure supplement 4***. ***Figure 3—source data 1*** shows the normalized enrichmentscores of the hallmarks identified in ***Figure 3—figure supplement 6***. ***Figure 3—source data 2*** shows the Ct values from the qRT-PCR analysis in ***Figure 1—figure supplement 4*** and ***Figure 3— figure supplement 7***. ***Figure 4—source data 1*** shows the full-length gel images of recombination analysis in ***Figure 4—figure supplement 2***.

***Supplementary file 1*** shows the gel densitometer results of recombination analysis of 14 organs after tamoxifen injection in ***Figure 4—figure supplement 2***. ***Supplementary file 2*** shows the statistical analysis of pairwise comparisons of the survival plots of the four *LSL-Kras* alleles in ***Figure 4A***. ***Supplementary file 3*** shows the histopathology report for H&E stained tissues from multiple organs from each of the four *LSL-Kras* alleles. ***Supplementary file 4*** shows the grading of observed tumors in multiple organs from each of the four *LSL-Kras* alleles.

## Acknowledgements

We thank members of the Counter Laboratory for technical assistance, David Kirsch, James Alvarez, Mark Onaitis, Channing Der, and Aaron Hobbs for advice, Cheryl Block, Gary Kucera, and the Duke Cancer Institute Transgenic Mouse Facility for the generation of mice, and Nicolas Devos, David Corcoran, Wei Chen, and Duke Center for Genomic and Computational Biology for their assistance in RNA-Seq.

This work was supported by the National Cancer Institute (R01CA94184 and P01CA203657 to CMC) and the Shared Resources of the Duke Cancer Institute (P30CA014236). The authors declare no competing interests.

## Author Contributions

O. Erdogan, N.L.K. Pershing, E. Kaltenbrun, and C.M. Counter conceived the study. O. Erdogan and C.M. Counter designed the experiments and the methodology. O. Erdogan and E. Kaltenbrun established the mouse models. O. Erdogan performed the tumor studies, cell culture experiments, RAS activity assays, tissue isolation, RNA extraction, and RNASeq. N.J. Newman performed recombination PCR. O. Erdogan analyzed recombination PCR data, tumor studies, and transcriptome data. J.I. Everitt analyzed the histopathology. C.M. Counter supervised the study. O. Erdogan, N.L.K. Pershing, J.I. Everitt, and C.M. Counter wrote the manuscript. All authors read, edited, and agreed on the final version of the manuscript.

---

KEY RESOURCE TABLE
| REAGENT or RESOURCE | SOURCE | IDENTIFIER |
| --- | --- | --- |
| **Antibodies** |  |  |
| FLAG | Sigma | F1804 |
| KRAS | Santa Cruz Biotechnology | sc-30 |
| Beta-Tubulin | Sigma | T5201 |
| **Bacteria strains** |  |  |
| STBL3 | Thermo Fisher Scientific | C737303 |
| **Chemicals, peptides, recombinant proteins** |  |  |
| DMEM with high glucose | Sigma-Aldrich | D5796 |
| Fetal Bovine Serum | VWR | 97068-085 |
| FuGENE® 6 Transfection Reagent | Promega | E2691 |
| Protease inhibitor cocktail | Roche | 11836170001 |
| DC Protein Assay | Bio-Rad | 5000112 |
| Geneticin | Thermo-Fisher | 10131035 |
| Hygromycin | Thermo-Fisher | 10687010 |
| DNeasy Blood and Tissue DNA extraction Kit | Qiagen | 69504 |
| Platinum Taq polymerase | Invitrogen | 10342046 |
| Tamoxifen | Sigma-Aldrich | T5648-5G |
| Corn oil | Sigma-Aldrich | C8267 |
| RNaseZAP™ | Sigma-Aldrich | R2020 |
| RNA Later | Sigma-Aldrich | R0901 |
| RNAeasy Kit | Qiagen | 74104 |
| QIASHredder columns | Qiagen | 79654 |
| RNase-Free DNase Set | Qiagen | 79254 |
| RNAsecure™ RNase Inactivation Reagent | Thermo-Fisher | AM7005 |
| **Critical Commercial Assays** |  |  |
| Active Ras Detection Kit | Cell Signaling | 8821 |
| Ras GTPase ELISA Kit | Abcam | ab134640 |
| **Cell lines** |  |  |
| HEK-HT | (Counter et al., 1992) |  |
| **Mouse Strains** |  |  |
| LSL-Kras<sup>natG12D</sup>/+ | This study |  |
| LSL-Kras<sup>natQ61R</sup>/+ | This study |  |
| LSL-Kras<sup>comG12D</sup>/+ | This study |  |
| LSL-Kras<sup>comQ61R</sup>/+ | This study |  |
| ACTB:FLPe/+ | Jackson Laboratory | 003800 |
| CC10-CreER/+;R26-CAG-fGFP/+ | (Xu et al., 2012) |  |
| Rosa26-CreERT2/+ | Jackson Laboratory | 008463 |
| **Recombinant DNA** |  |  |
| pcDNA3.1+ FLAG-Kras<sup>natG12D</sup> | This study |  |
| pcDNA3.1+ FLAG-Kras<sup>natQ61R</sup> | This study |  |
| pcDNA3.1+ FLAG-Kras<sup>comG12D</sup> | This study |  |
| pcDNA3.1+ FLAG-Kras<sup>comQ61R</sup> | This study |  |
| pBABE-neo largeTcDNA | (Hahn et al., 2002) | Addgene #1780 |
| MSCV-Cre-Hygro | (Wang et al., 2010) | Addgene #34565 |
| **Software and algorithms** |  |  |
| ImageJ version 1.52k with Java 1.8.0_172 | NIH (Schneider et al., 2012) | https://imagej.nih.gov/ij/ |
| Image Lab | Bio-Rad | https://www.bio-rad.com/en-us/product/image-lab-software |
| Graphpad Prism v8 | Graphpad | https://www.graphpad.com/ |
| bcl2fastq | Illumina | https://support.illumina.com/sequencing/sequencing_software/bcl2fastq-conversion-software.html |
| STAR RNA-Seq alignment tool | (*Dobin et al., 2013*) | https://github.com/alexdobin/STAR |
| DESeq2 | (*Love et al., 2014*) | http://www.bioconductor.org/packages/release/bioc/html/DESeq2.html |
| GSEA | (*Mootha et al., 2003*) | https://www.gsea-msigdb.org/gsea/ |
| R Studio |  | https://www.R-project.org |

**Figure 1-figure supplement 1.**
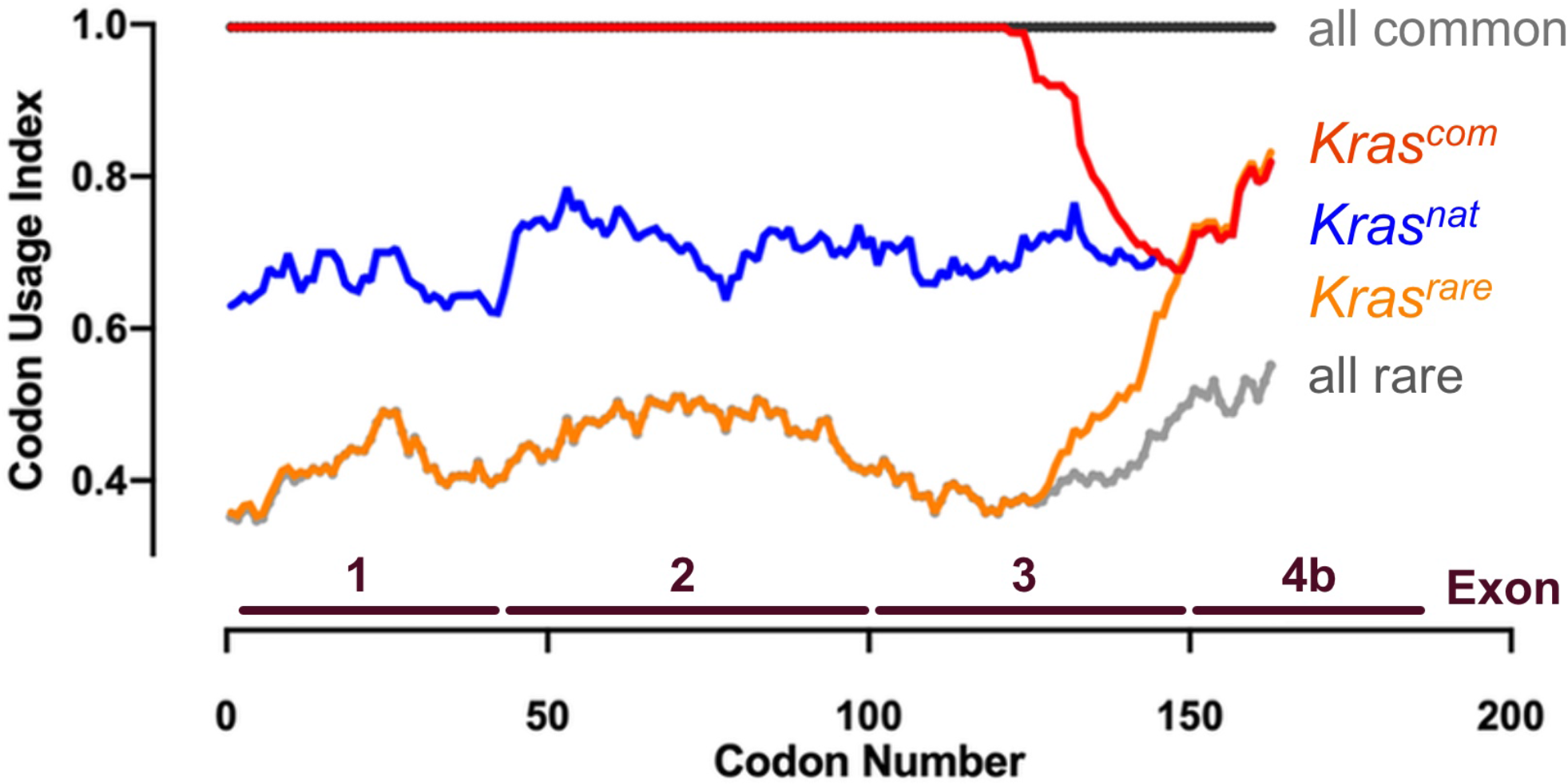
Codon usage of the four novel *LSL-Kras* alleles. Codon usage index versus codon number of *Kras* with native (nat, blue line), common (com, red line) and rare (rare, orange line) codons. Maximum rare (all rare, grey dotted line) and common (all common, solid grey line) codon usage are shown for reference.

**Figure 1-figure supplement 2.**
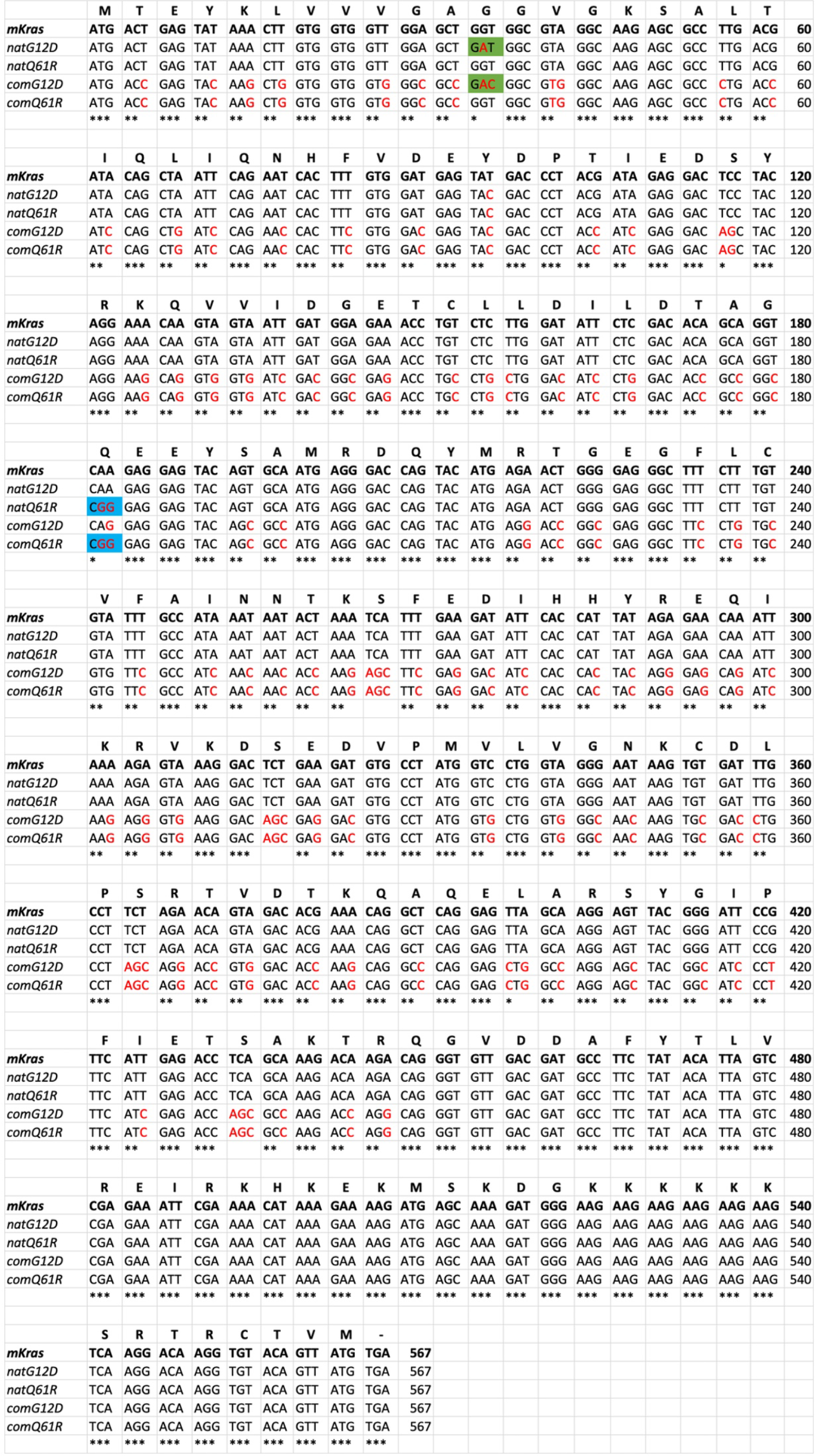
Coding sequence of exons 1 to 3 of the four novel *LSL-Kras* alleles. Sequence alignment of the codons in the coding exons 1-3 of the indicated *LSL-Kras* alleles in comparison to mouse *Kras* transcript **(mKras).** *Top:* amino acid sequence . *Red nucleotides:* optimized codons. *Green highlight:* G12D mutation. *Blue highlight:* 061R mutation.

**Figure 1-figure supplement 3.**
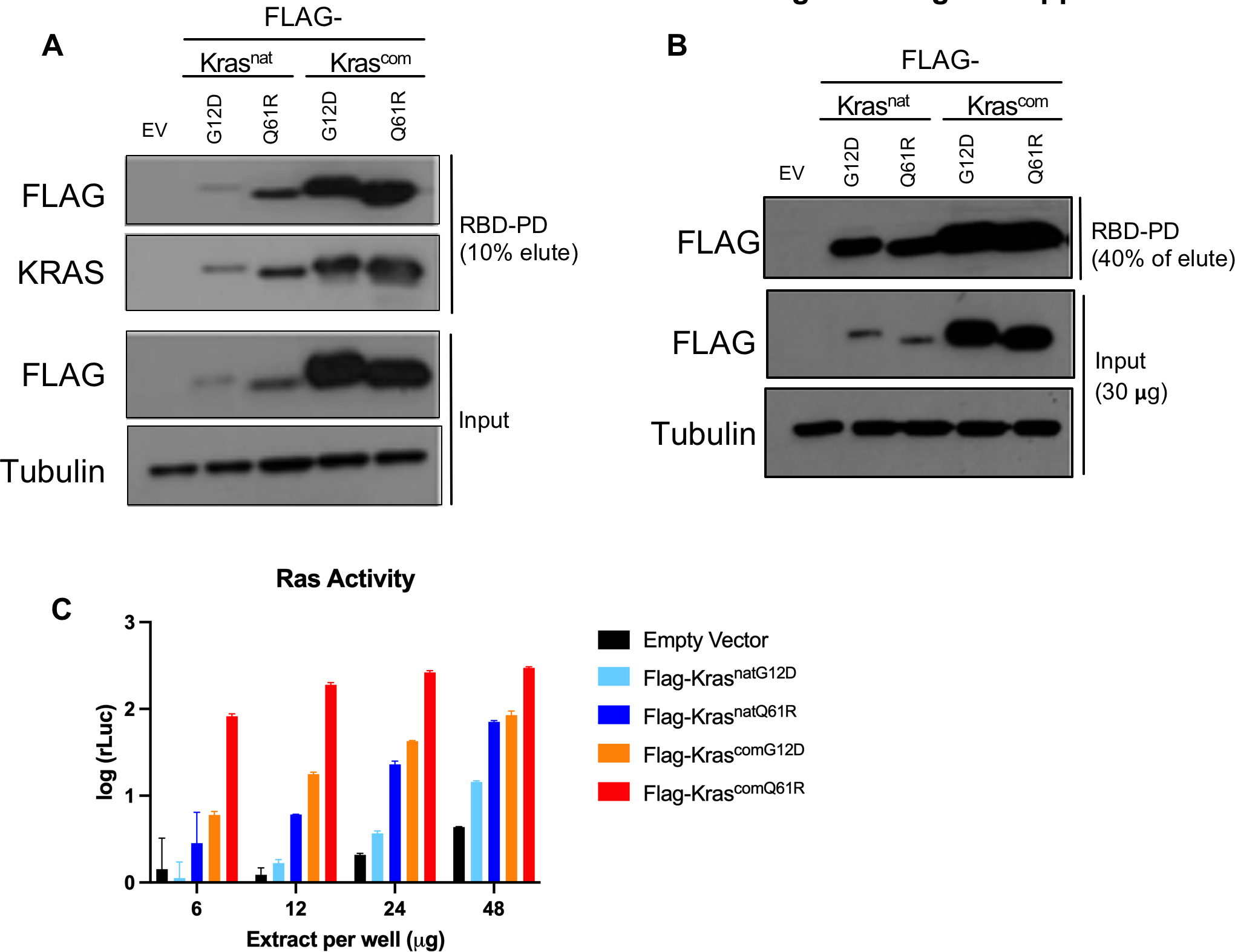
Expression and activity level of the four novel encoded *Kras* proteins. Expression levels, determined by immunoblot with an anti-FLAG antibody, and activity levels, determined by RBD pull-down (RBD-PD) followed by (A,B) immunoblots of lysates derived from HEK-HT cells transiently expressing the indicated FLAG-tagged Kras proteins and (C) by ELISA as a biological replicate of Figure 1C. EV: Empty Vector . A and B show two biological replicates. Tubulin and empty vector serve as loading and negative controls, respectively. Full-length gel images are provided at Figure 1*-source data* 2.

**Figure 1-figure supplement 4.**
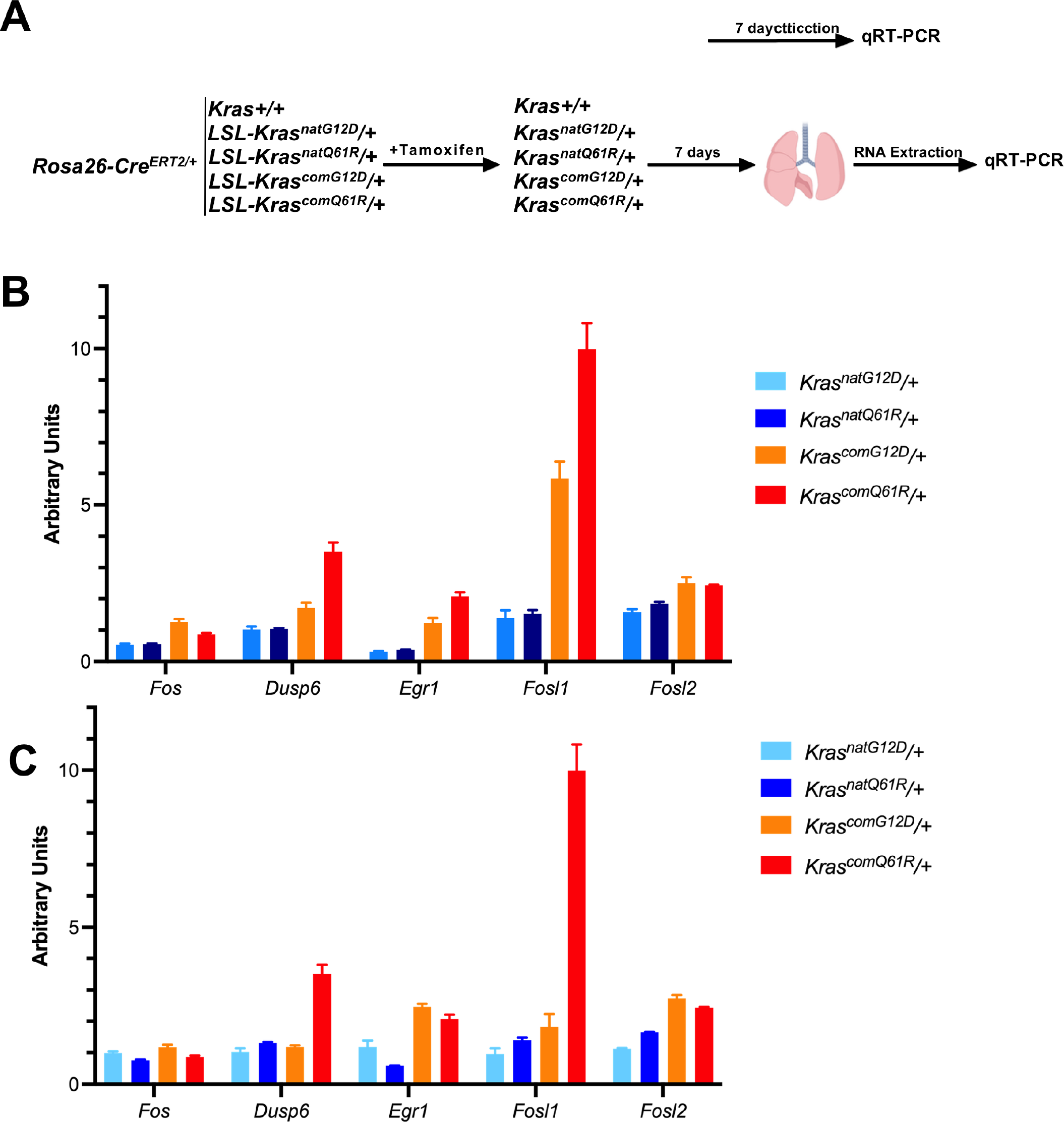
qRT-PCR validates an increase in RAS target gene expression with increased Kras activity. 2 mice from cohorts of *LSL-KrasnatG ^120^!+, LSL-Krasnat0 ^61^R/+, LSL-KrascomG ^120^!+,* and *LSL-Krascom0 ^61^R/+* in a *Rosa26-CreERT2/+* background, injected with tamoxifen at 6 to 8 weeks of age and RNA was extracted from lungs seven days later. 2 *Rosa26-CreERT2/+* mice were treated similarly as a control (A). First strand cDNA synthesis and qRT-PCR of selected genes downstream ERK/MAPK pathway was performed using GoTaq 2-Step RT-qPCR kit (B,C). All measurements were normalized against Actin as the internal control using the 2-LILICt method. B and C are biological replicates.

**Figure 2-figure supplement 1.**
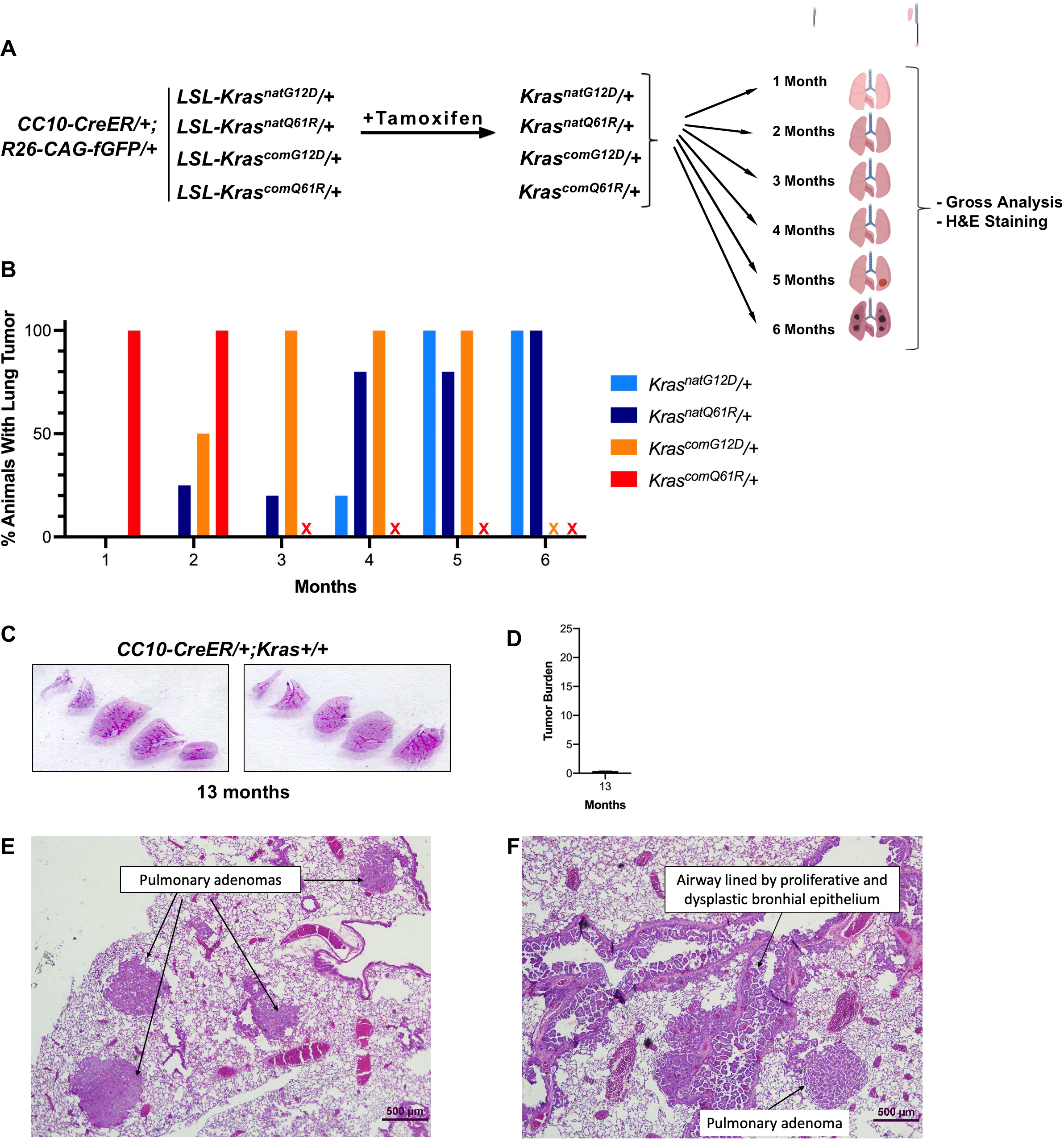
Tumor phenotypes track with intrinsic oncogenic Kras signaling. (A) Schematic of the experimental design. (B) % of mice with at least one visible lung lesion. (C) Representative H&E staining of lungs and (D) mean ± SD % tumor burden from microscopic analysis of two lung sections from two *CC10-CreERl+;Kras+!+* mice 13 months after tamoxifen . (E,F) Predominant proliferative lesions in lungs of *CC10-CreERl+;KrascomG ^120^1+* and *CC10-CreERl+;KrascomQ ^61^R;+* differ. Photomicrograph of lung section from (E) *CC10-CreER!+;KrascomG ^120^!+* mouse at 4 months showing multiple large pulmonary adenomas, and (F) *CC10-CreER!+;KrascomQ ^61^R;+* mouse at 1 month showing extensive proliferation of airway epithelium into papillary projections of hyperplastic and dysplastic cells as well as pulmonary adenoma. H&E staining, x40 magnification .

**Figure 3-figure supplement 1.**
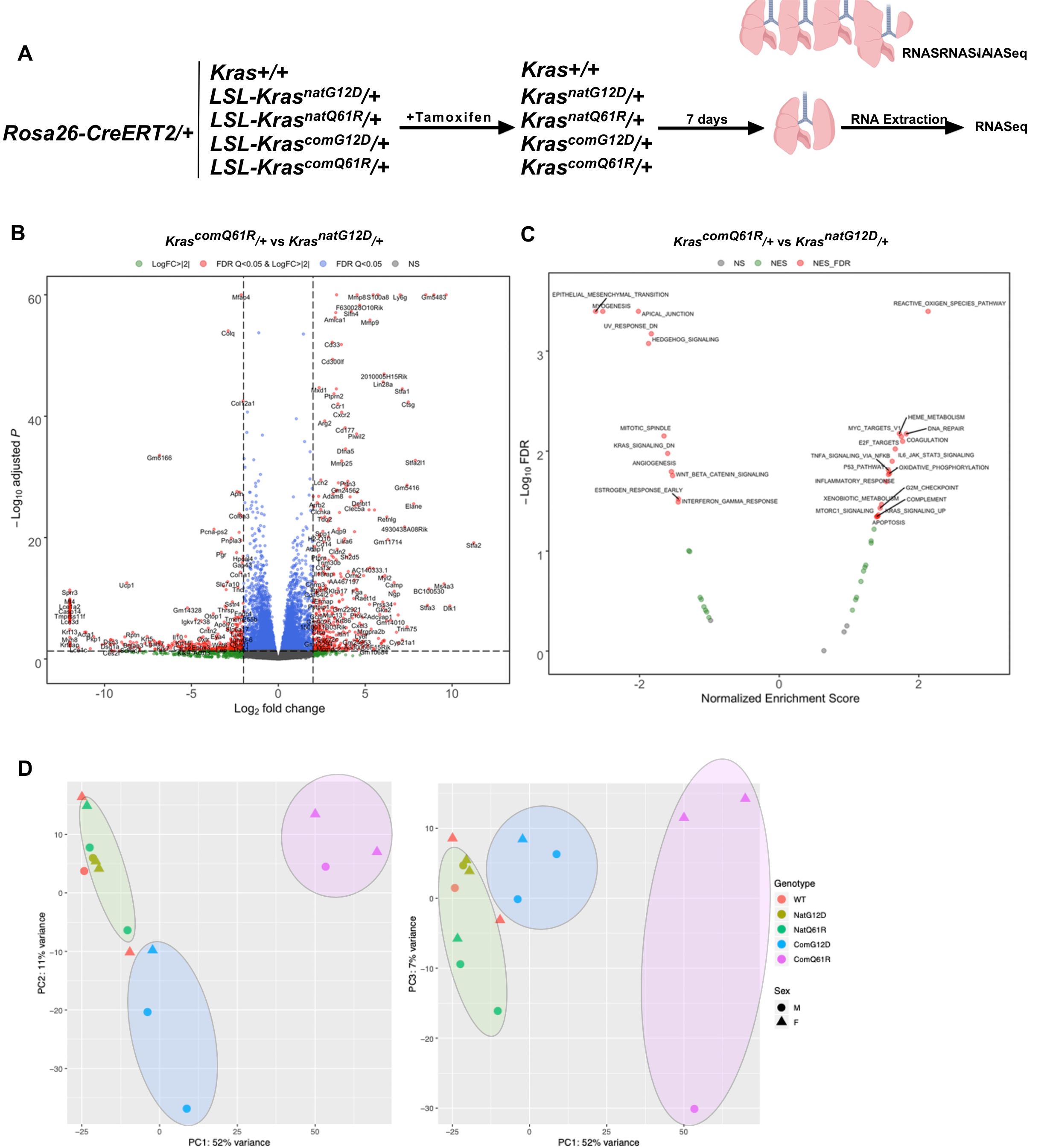
Transcriptomes of each oncogenic *LSL-Kras* allele. **(A)** Schematic of the experimental design. **(B)** Volcano plot of the */og2* fold-change versus *p* value of differentially expressed genes and **(C)** the false discovery rate (FDR) versus normalized enrichment score of the indicated GSEA hallmarks determined from RNA-seq analysis of the lungs of *Rosa26- CreERT2/+;LSL-Krascomo51R;+* versus *Rosa26-CreERT2/+;LSL-Krasn atG ^120^!+* mice seven days after tamoxifen injection. **(D)** Principal component analysis of the transcriptome signatures of engineered alleles . Alleles with high quantitative signaling are highlighted with blue *(KrascomG ^120^!+)* and pink shade *(Krascomo^51^RJ+),* and alleles with and low signaling *(Krasnat)* in green shade.

**Figure 3-figure supplement 2.**
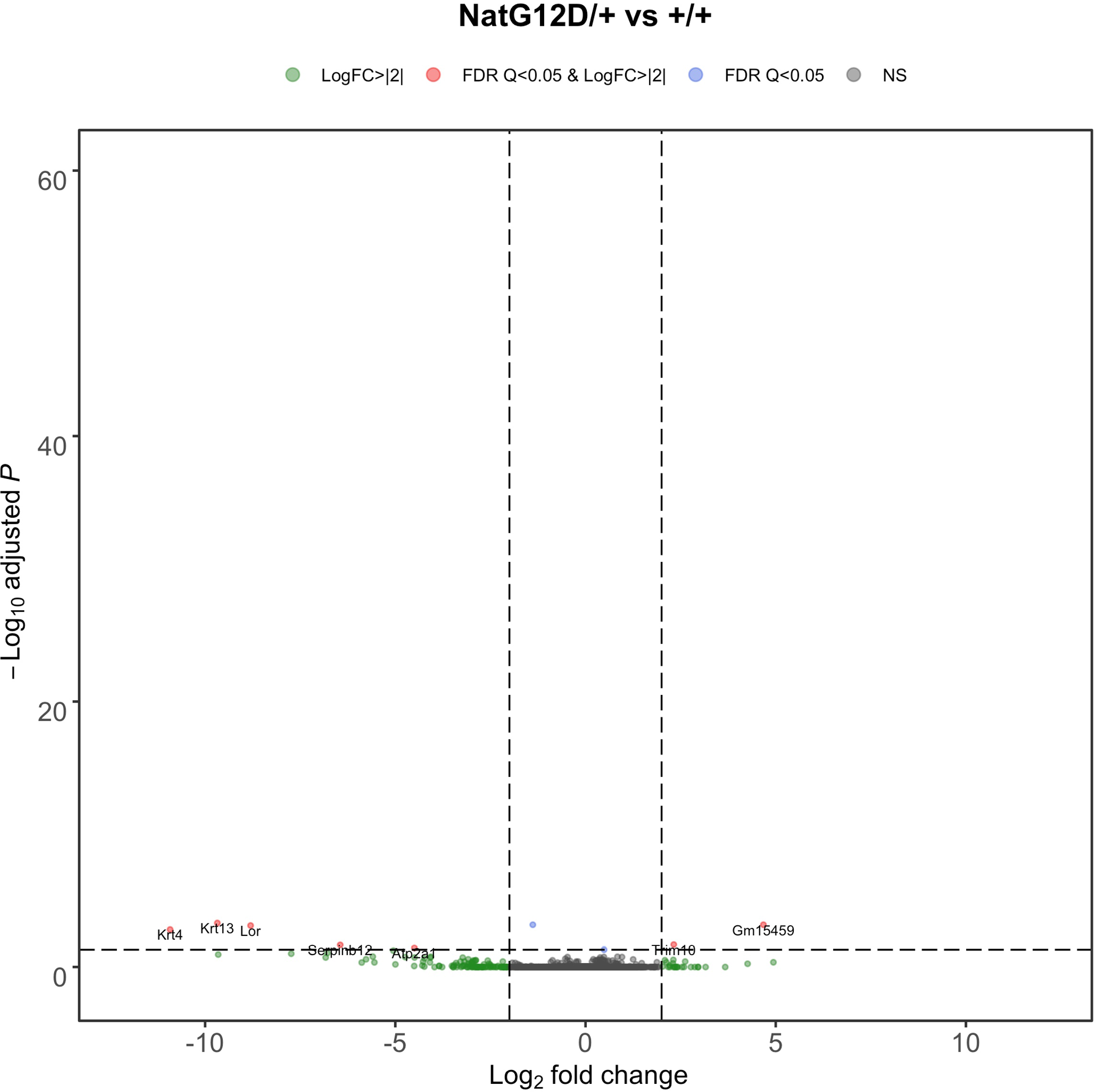
Transcriptome upon activating the *LSL-KrasnatG ^120^/+* allele. Volcano plot of the */og2* fold-change versus *p* value of differentially expressed genes from RNA-seq analysis of the lungs of *Rosa26- CreERT2/+;LSL-KrasnatG120!+* versus *Rosa26-CreERT2/+;Kras+/+* mice seven days after tamoxifen injection .

**Figure 3-figure supplement 3.**
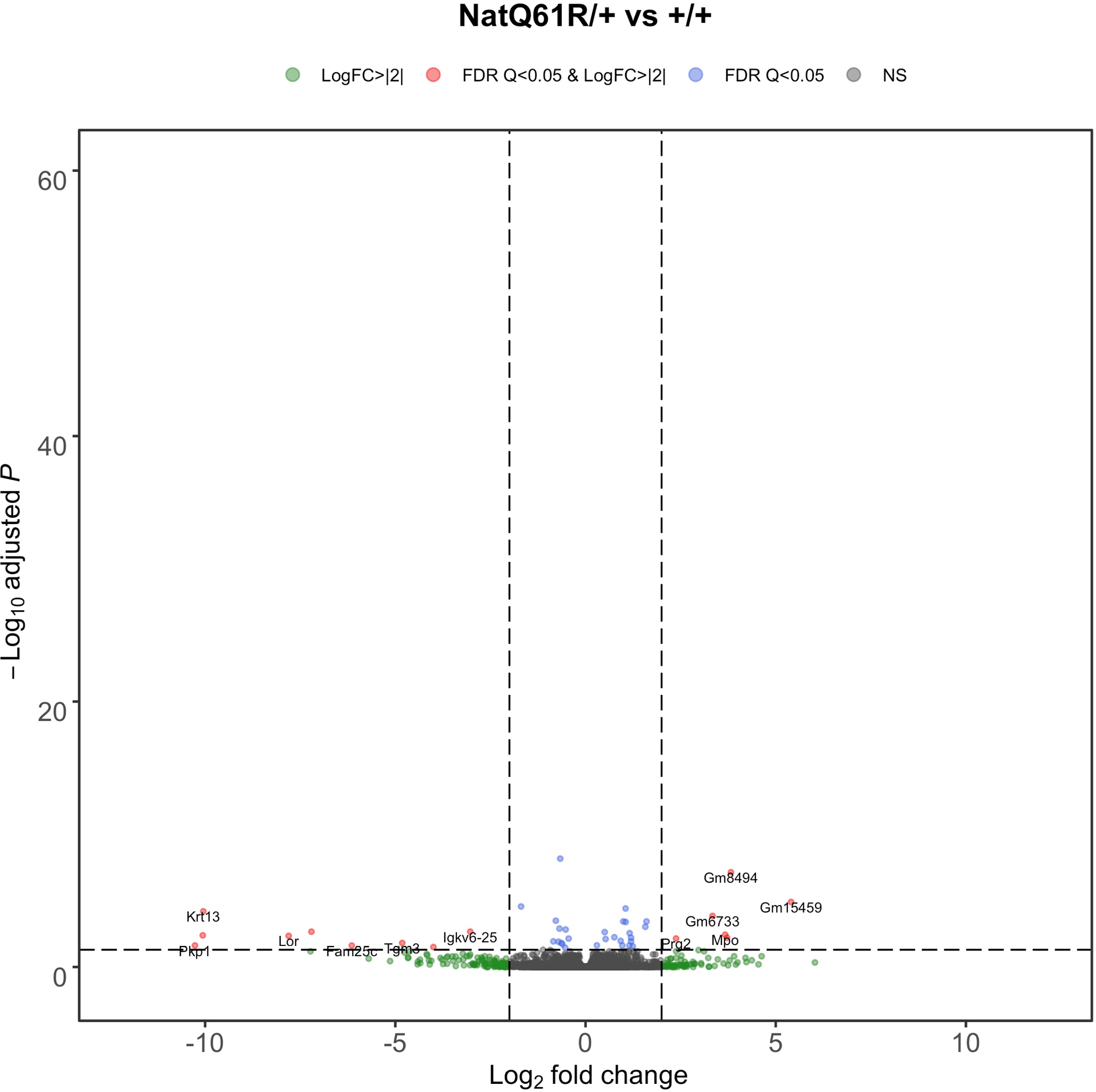
Transcriptome upon activation of the *LSL-KrasnatQGf Rj+* allele. Volcano plot of the */og2* fold-change versus *p* value of differentially expressed genes from RNA-seq analysis of the lungs of *Rosa26- CreERT2/+;LSL-Krasnat061R/+* versus *Rosa26-CreERT2/+;Kras+/+* mice seven days after tamoxifen injection .

**Figure 3-figure supplement 4.**
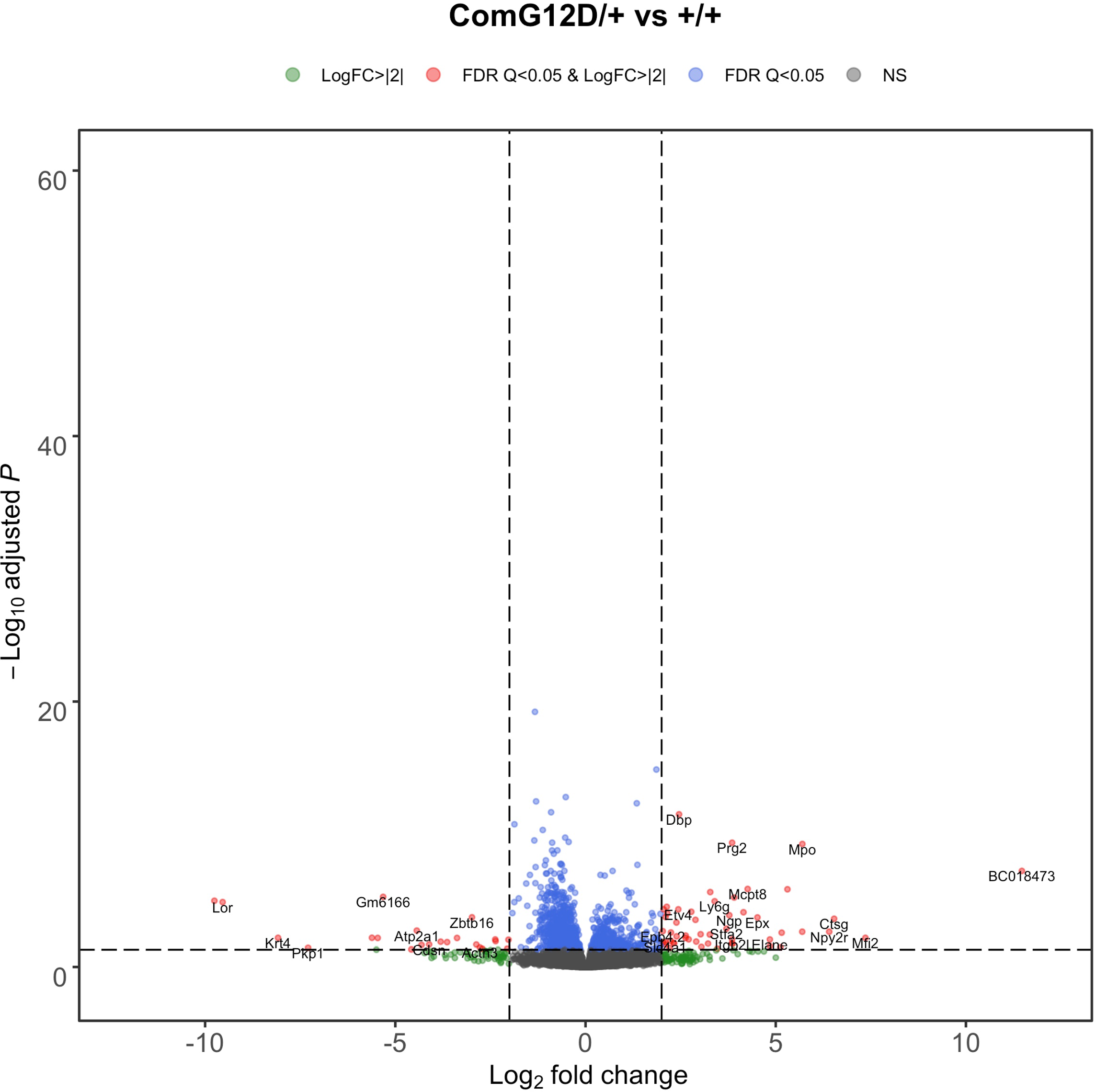
Transcriptome upon activating the *LSL-KrascomG ^120^/+* allele. Volcano plot of the */og2* fold-change versus *p* value of differentially expressed genes from RNA-seq analysis of the lungs of *Rosa26- CreERT2/+;LSL-KrascomGt2D;+* versus *Rosa26-CreERT2/+;Kras+/+* mice seven days after tamoxifen injection.

**Figure 3-figure supplement 5.**
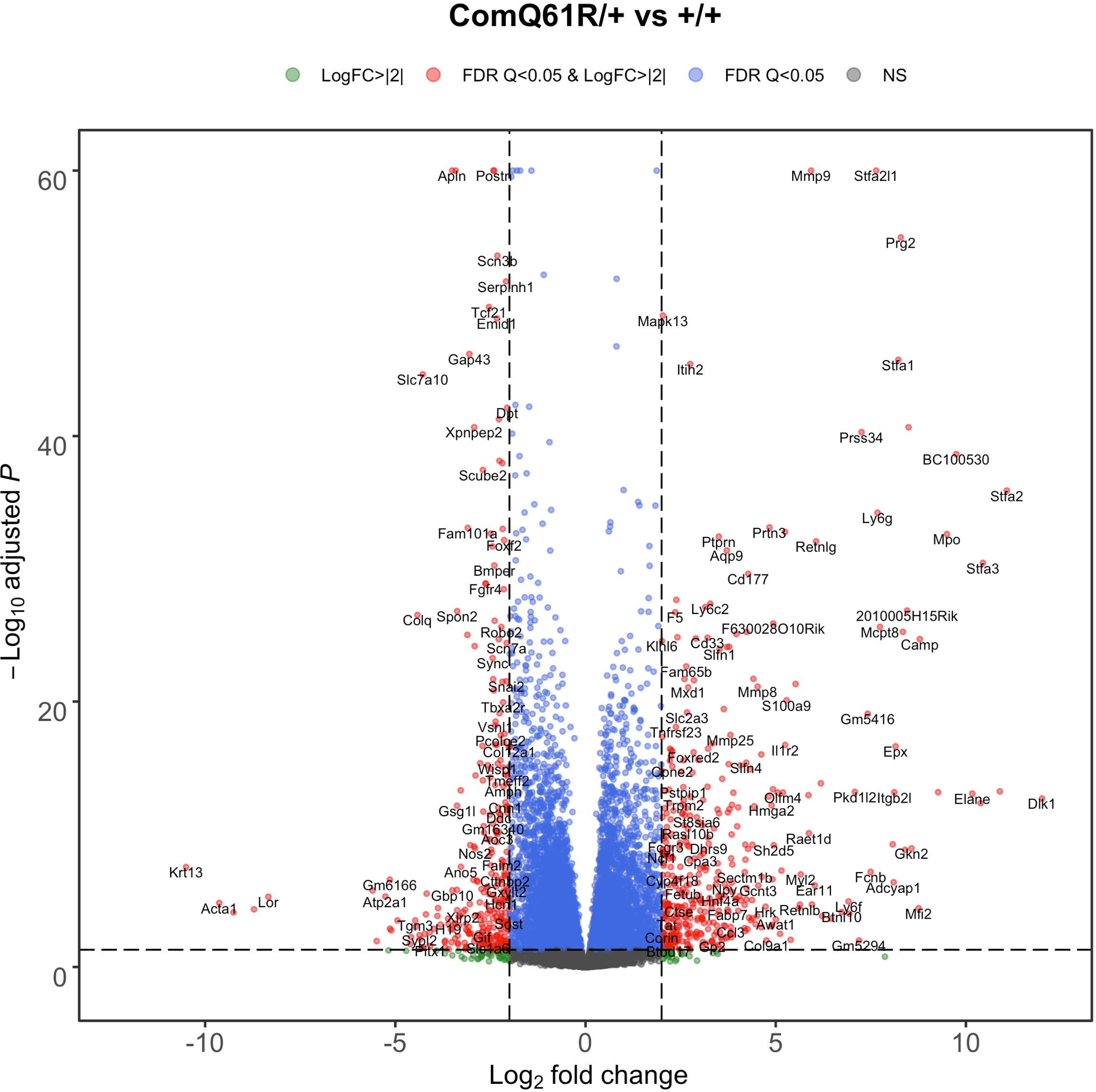
Transcriptome upon activating the *LSL-KrascomQGf Rj+* allele. Volcano plot of the */og2* fold-change versus *p* value of differentially expressed genes from RNA-seq analysis of the lungs of *Rosa26- CreERT2/+;LSL-KrascomOB 1R/ +* versus *Rosa26-CreERT2/+;Kras+/ +* mice seven days after tamoxifen injection.

**Figure 3-figure supplement 6.**
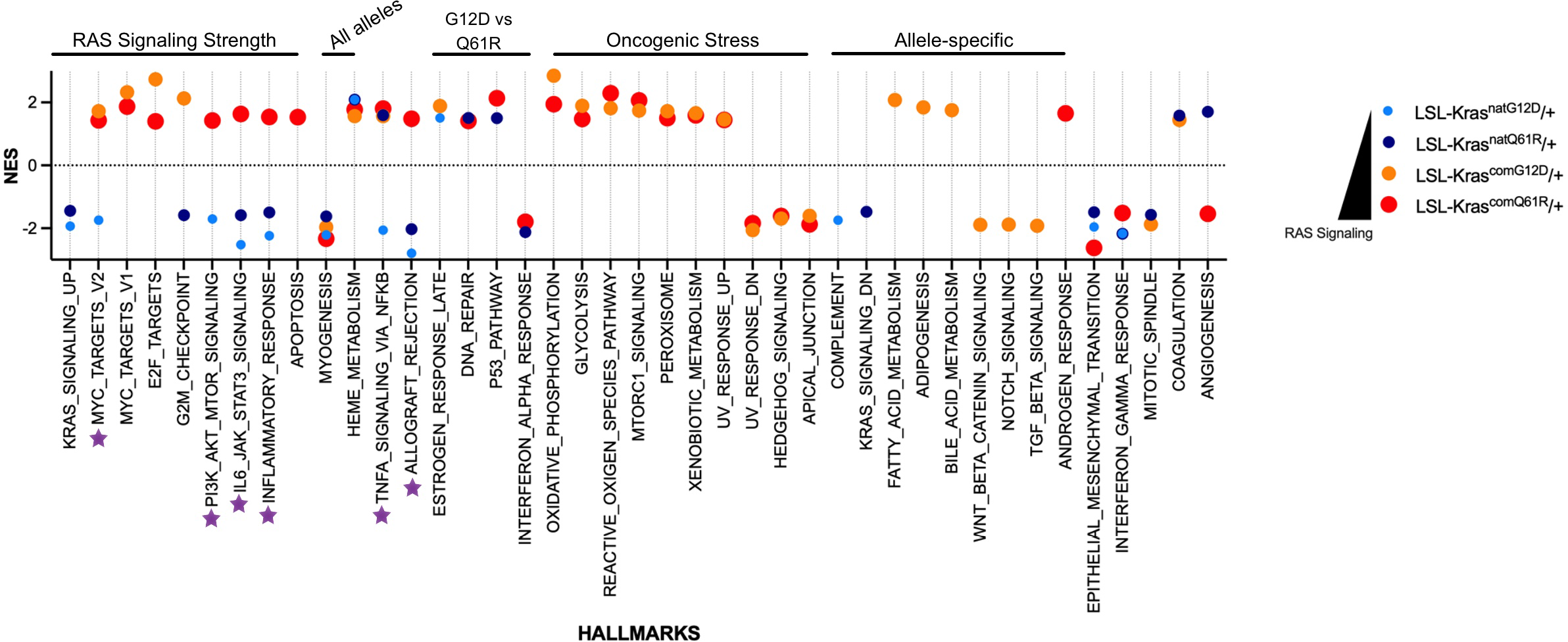
Transcriptome GSEA hallmarks upon activating the panel of *LSL-Kras* alleles. Purple stars show alleles that show qualitative signaling dependent enrichment or depletion patterns of hallmarks. Only hallmarks with **FDR** less than 5% are shown. Dot size is adjusted according to RAS activity for better visualization .

**Figure 3-figure supplement 7.**
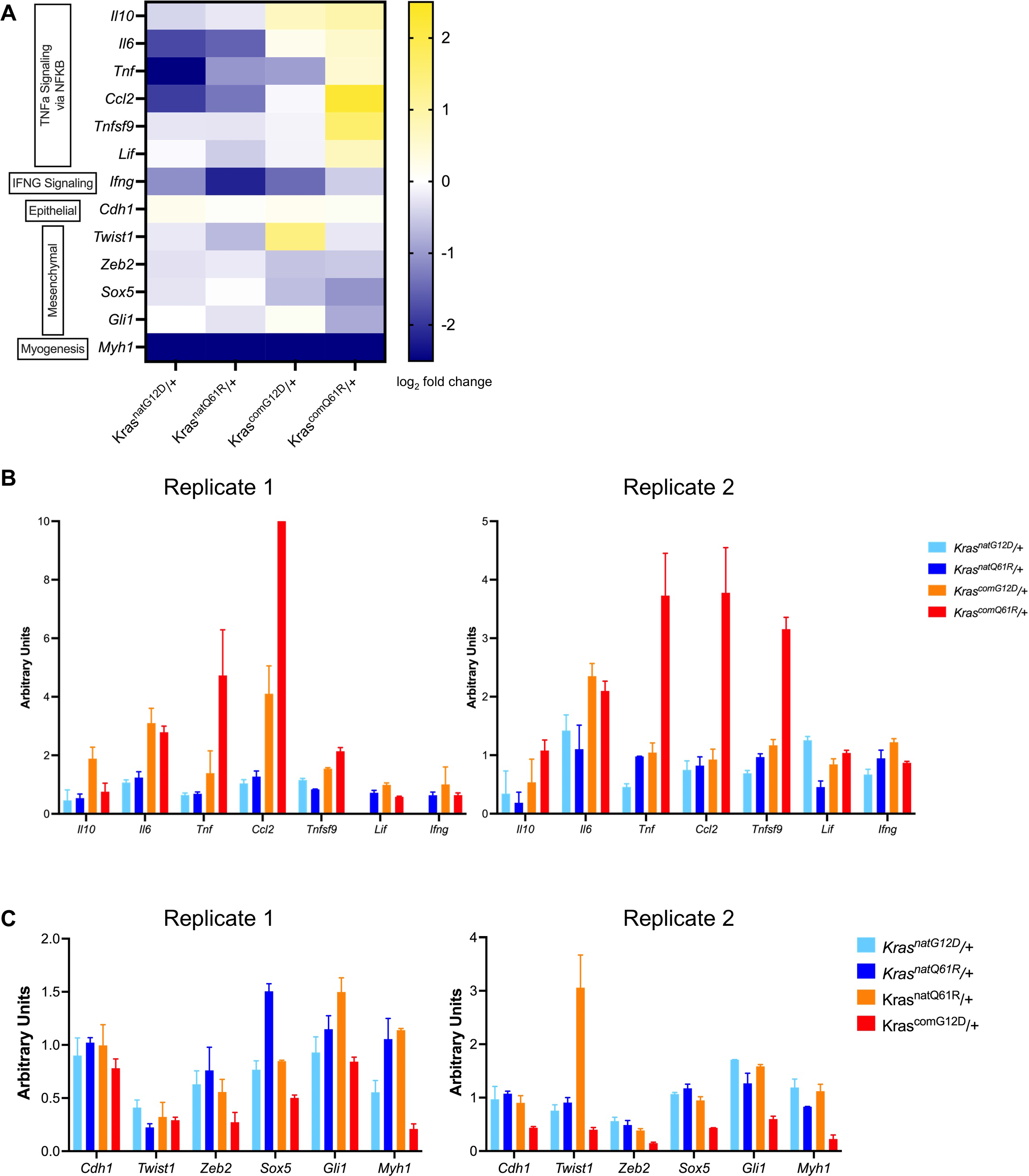
Validation of signaling responses via **qRT-PCR** analysis of the selected genes. **(A)** heatmap of the log_2_ fold-change of selected genes from RNAseq analysis of the lungs of indicated *LSL-Kras* alleles in the *Rosa26- CreERT2/+* background in comparison to the wild type *Kras* allele seven days after tamoxifen injection. (B,C) Validation of signaling responses via qRT-PCR analysis of the selected genes in **A** from pathways of TNFa Signaling via NF-KB and lnterferony Signaling (B),and EMT and Myogenesis (C). Two biological replicates.

**Figure 4-figure supplement 1.**
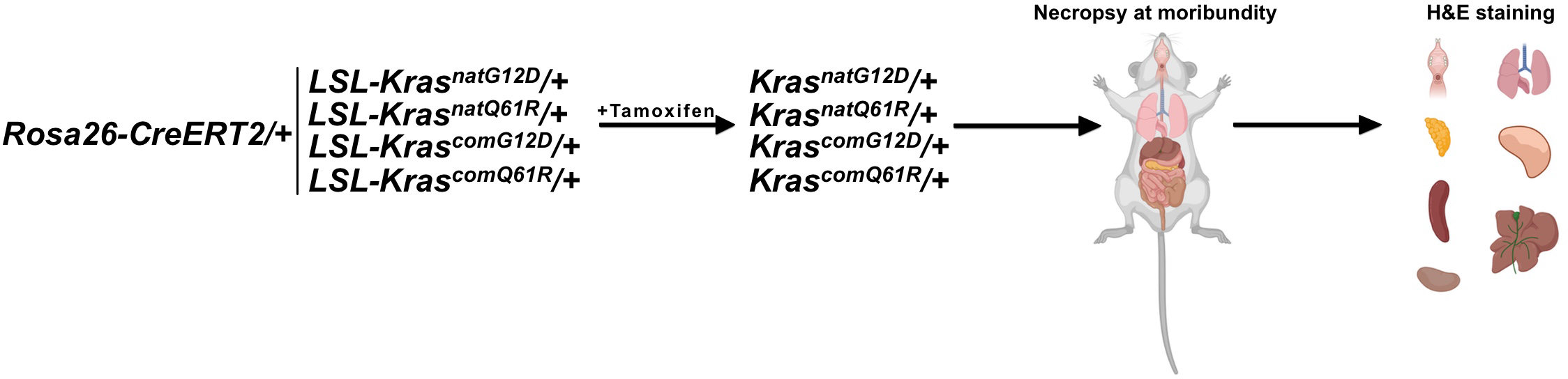
Global activation of each engineered *LSL-Kras* allele using a ubiquitous Cre driver. Schematic of the experimental design. Cohorts of three adult male and female *Rosa26-CreERT2/+;LSL-KrasnatGt2D/+, Rosa26- CreERT2/+;LSL-KrasnatOBtR/+, Rosa26-CreERT2/+;LSL-KrascamGt2D/+,* and *Rosa26-CreERT2/+;LSL-KrascamQBtR/+* mice, and as a control for normalization, *Rosa26-CreERT2/+;Kras+/+ mice,* were injected with tamoxifen and humanely euthanized at moribundity . Selected set of organs were fixed in formalin, sliced and embedded in paraffin, sectioned, and **H&E** stained.

**Figure 4-figure supplement 2.**
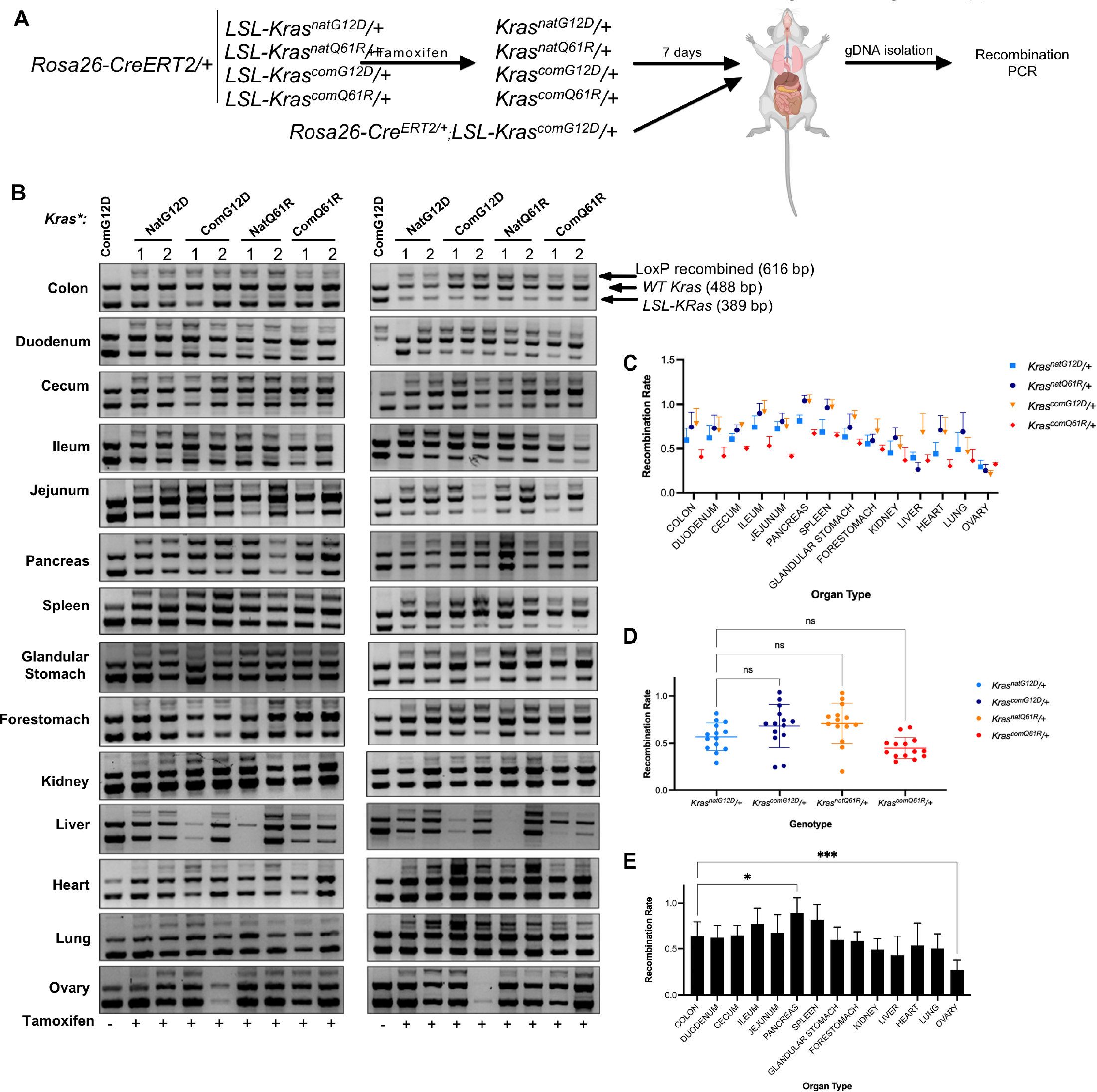
Tissue-specific recombination rates of each oncogenic *LSL-Kras* allele upon global activation. (A) Schematic of the experimental design. Two mice from each *LSL-Kras* genotype in a *Rosa26-CreERT2/+* background 7 days after tamoxifen injection were humanely euthanized, and their organs were removed. Genomic DNA was isolated and subjected to genotyping PCR. One mouse in *Rosa26-CreERT2/+ ; LSL-KrascomG ^120^!+* background was used as a no tamoxifen control. (B) Gel images from duplicate recombination analysis and (C) mean ± SEM of the ratio of the recombined *LSL-Kras* (LoxP recombined, 616 bp) to unaltered wild-type *Kras* (WT, 488 bp) allele products determined by PCR genotyping. Two technical replicates of two biological replicates . Full-length gel images are provided at Figure 4*-source data 1*. (D,E) One-way Anova with Bonferroni’s multiple comparisons test with a single pooled variance and a 95% Cl shows variation in recombination rate across different alleles (D) compared to the *KrasnatG ^120^!+* allele or across organs (E) compared to the colon, with the exception of ovary, was not statistically significant (*p<0.1 and ***p<0 .001).

**Figure 4-figure supplement 3.**
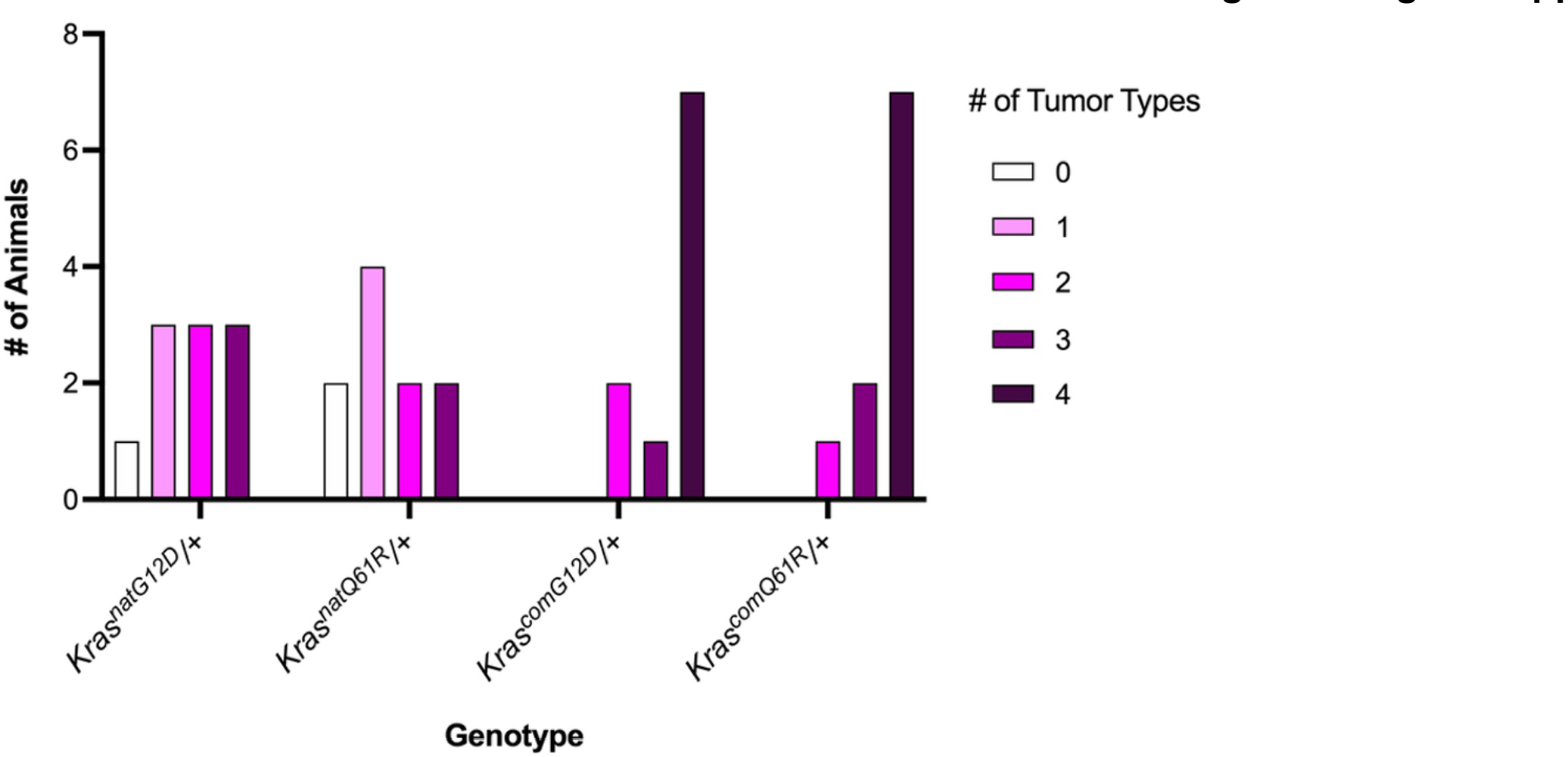
Number of tumor types induced upon globally activating each oncogenic *LSL-Kras* allele. Number of mice with the indicated number of different tumor types are shown (see *Supplementary File* 3).

**Figure 4-figure supplement 4.**
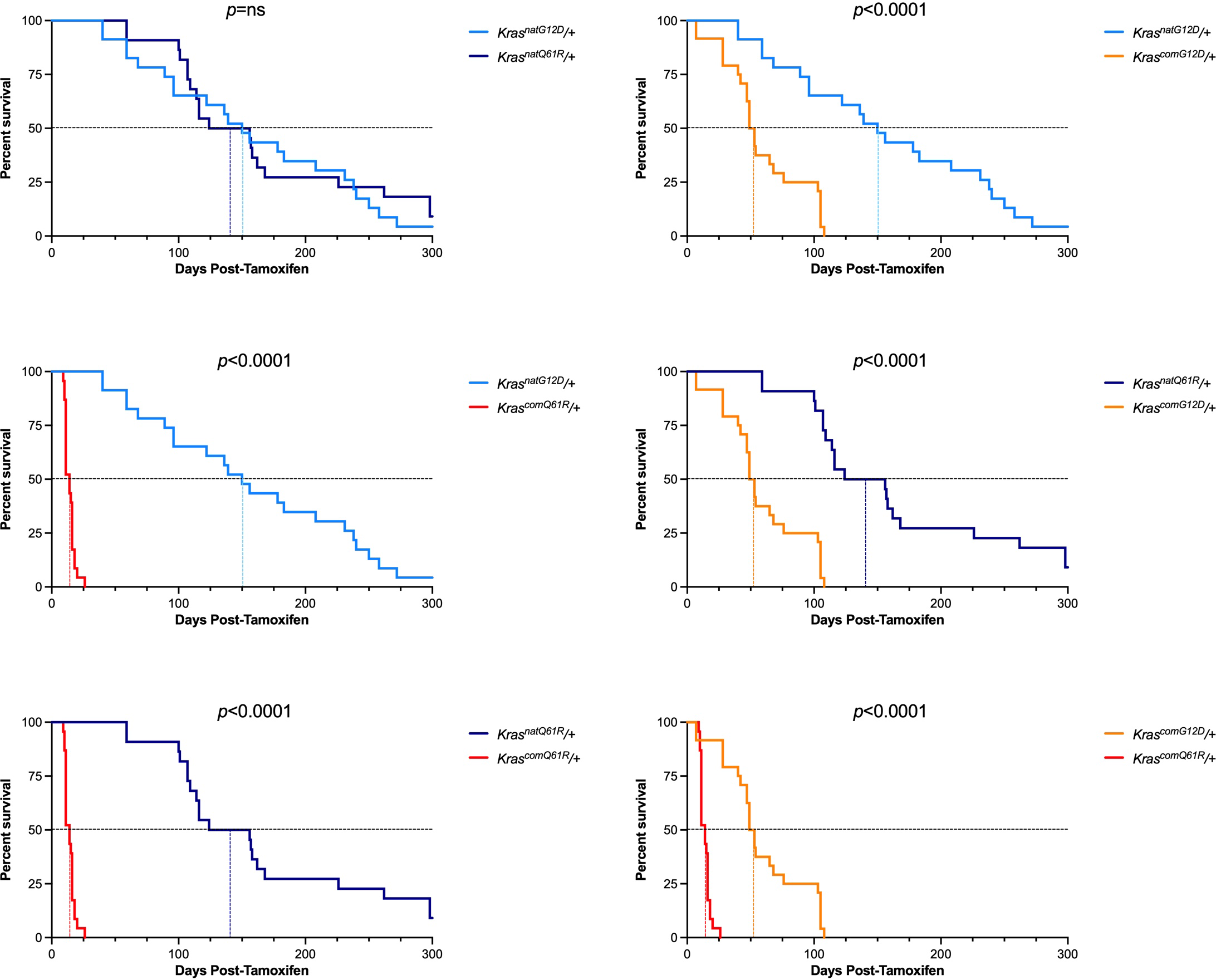
Pairwise comparison of the survival curves of the engineered *LSL-Kras* alleles upon global activation in the *Rosa26-CreERT2/+* background mice after tamoxifen injection. *p* values calculated by Log-Rank (Mantel-Cox) test of the pairwise comparisons of alleles (see *Supplementary File* 2).

**Figure 4-figure supplement 5.**
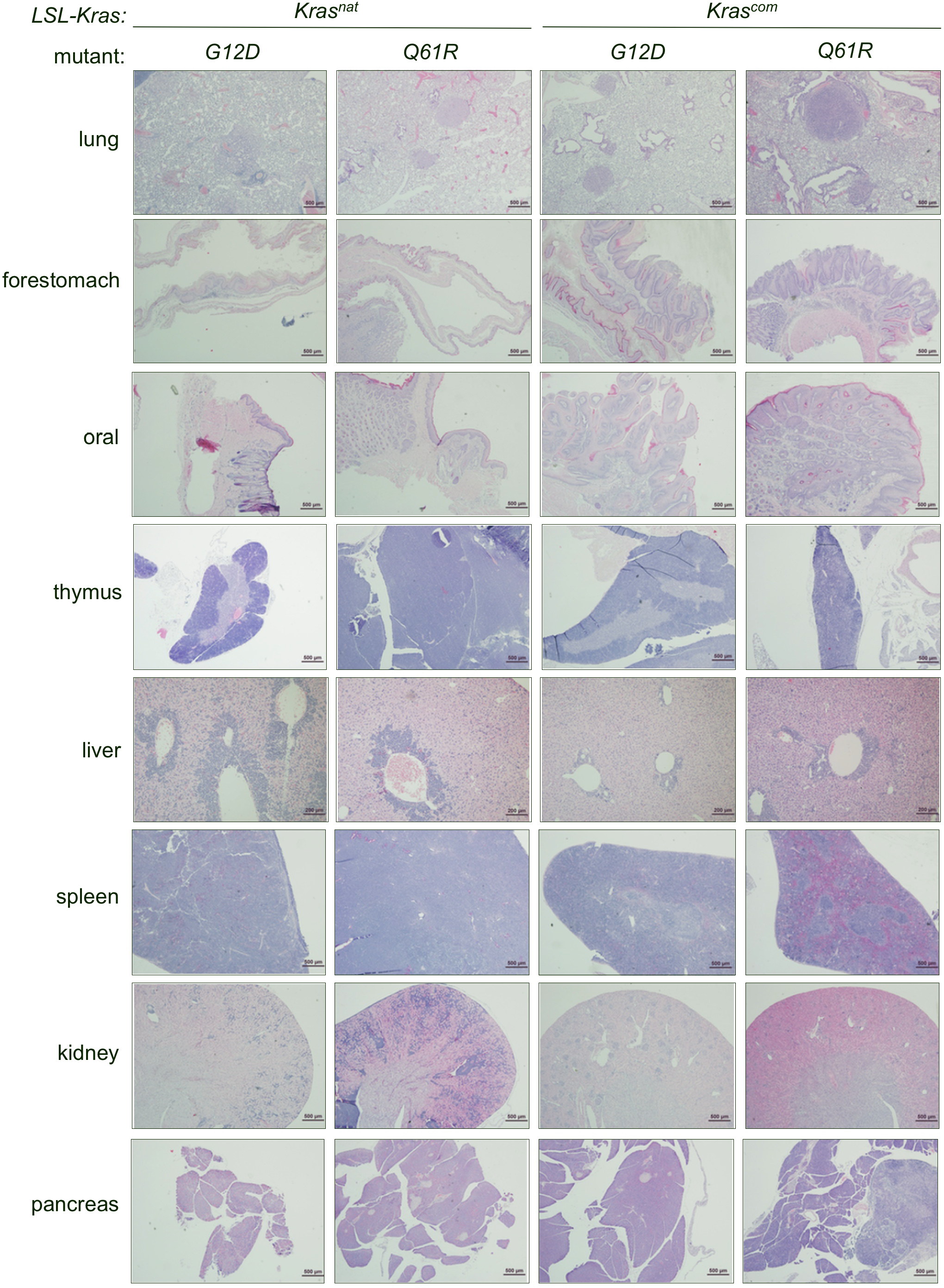
Proliferative lesions induced upon activating each oncogenic *LSL-Kras* allele. Examples of H&E stained sections from the indicated organs in each allele upon global activation at the moribundity endpoint.

**Figure 4-figure supplement 6.**
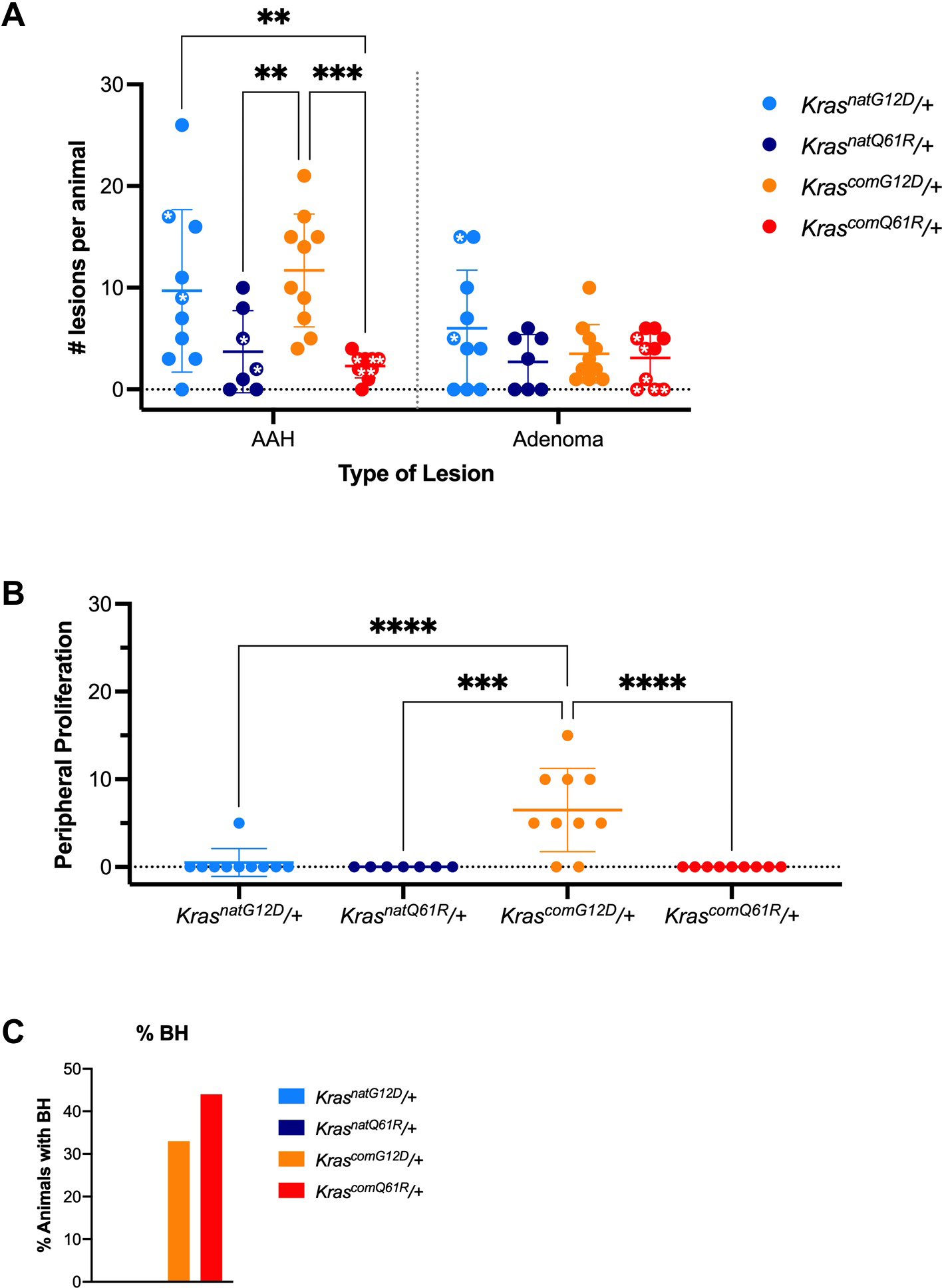
Histopathological analysis of pulmonary lesions induced upon activating each of the oncogenic *LSL-Kras* alleles. Mean ± SD of number of AHA and adenoma lesions per animal with stars denoting animals with extensive myeloid infiltration in the lungs **(A),** mean ± SD of peripheral lesions observed as percent surface area **(B),** and percent animals manifesting hyperplasia localized to bronchioles **(C).** *p* values calculated by two-way ANOVA multiple comparisons with Sidak testing **(A)** or one-way ANOVA multiple comparison with Tukey testing **(B).** ** p<0.01, *** p<0.001, and **** p<0.0001

**Figure 4-figure supplement 7.**
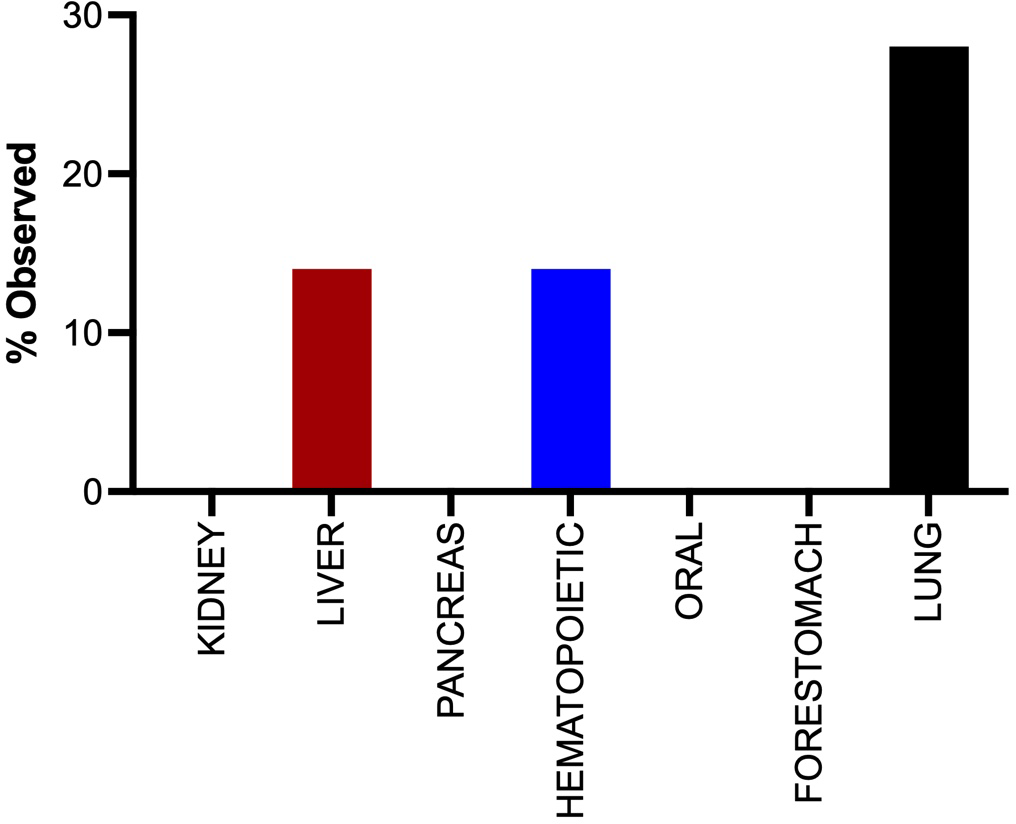
The tumor landscape of *Rosa26-CreERT2/+;KrasrareG ^120^/+* mice after tamoxifen injection. Percent of the indicated organs with tumors from *Rosa26-CreERT2/+;LSL-KrasrareG t2D/+* mice (n=7) 20 months after tamoxifen injection.

## References

1. Amendola CR, Mahaffey JP, Parker SJ, Ahearn IM, Chen WC, Zhou M, Court H, Shi J, Mendoza SL, Morten MJ, Rothenberg E, Gottlieb E, Wadghiri YZ, Possemato R, Hubbard SR, Balmain A, Kimmelman AC, Philips MR. 2019. KRAS4A directly regulates hexokinase 1. Nature 576: 482–486. DOI: https://doi.org/10.1038/s41586-019-1832-9, PMID: 31827279

2. Behringer R, Gertsenstein M, Nagy KV, Nagy A. 2014. Manipulating the mouse embryo: A laboratory manual (4^th^ edition). Cold Spring Harbor, New York. Cold Spring Harbor Laboratory Press.

3. Bos JL. 1989. *Ras* oncogenes in cancer: a review. Cancer Research 49: 4682–4689. PMID: 2547513

4. Buhrman G, Holzapfel G, Fetics S, Mattos C. 2010. Allosteric modulation of Ras positions Q61 for a direct role in catalysis. PNAS 107: 4931–4936. DOI: https://doi.org/10.1073/pnas0912226107, PMID: 20194776

5. Burd CE, Liu W, Huynh MV, Waqas MA, Gillahan JE, Clark KS, Fu K, Martin BL, Jeck WR, Souroullas GP, Darr DB, Zedek DC, Miley MJ, Baguley BC, Campbell SL, Sharpless NE. 2014. Mutation-specific RAS oncogenicity explains NRAS codon 61 selection in melanoma. Cancer Discovery 4: 1418–1429. DOI: https://doi.org/10.1158/2159-8290.CD-14-0729, PMID: 25252692

6. Burgess MR, Hwang E, Mroue R, Bielski CM, Wandler AM, Huang BJ, Firestone AJ, Young A, Lacap JA, Crocker L, Asthana S, Davis EM, Xu J, Akagi K, Le Beau MM, Li Q, Haley B, Stokoe D, Sampath D, Taylor BS et al., 2017. *KRAS* allelic imbalance enhances fitness and modulates MAP kinase dependence in cancer. Cell 168: 817–829.e15. DOI: https://doi.org/10.1016/j.cell.2017.01.020, PMID: 28215705

7. Cicchini M, Buza EL, Sagal KM, Gudiel AA, Durham AC, Feldser DM. 2017. Context-dependent effects of amplified MAPK signaling during lung adenocarcinoma initiation and progression. Cell Reports 18: 1958–1969. DOI: https://doi.org/10.1016/j.celrep.2017.01.069, PMID: 28228261

8. Counter CM, Avilion AA, LeFeuvre C, Stewart NG, Greider CW, Harley CB, Bacchetti S. 1992. Telomere shortening associated with chromosome instability is arrested in immortal cells which express telomerase activity. The EMBO Journal 11: 1921–1929. DOI https://doi.org/10.1002/j.1460-2075.1992.tb05245.x, PMID: 1582420

9. Dobin A, Davis CA, Schlesinger F, Drenkow J, Zaleski C, Jha S, Batut P, Chaisson M, Gingeras TR. 2013. STAR: ultrafast universal RNA-seq aligner. Bioinformatics 29: 15–21. DOI: https://doi.org/10.1093/bioinformatics/bts635, PMID: 2310488

10. Dymecki SM. 1996. Flp recombinase promotes site-specific DNA recombination in embryonic stem cells and transgenic mice. PNAS 93: 6191–6196. DOI: https://doi.org/10.1073/pnas.93.12.6191, PMID: 8650242

11. Edgar R, Domrachev M, Lash AE. 2002. Gene Expression Omnibus: NCBI gene expression and hybridization array data repository. Nucleic Acids Research 30: 207–210. DOI: https://doi.org/10.1093/nar/30.1.207, PMID: 11752295

12. Feil R, Brocard J, Mascrez B, LeMeur M, Metzger D, Chambon P. 1996. Ligand-activated site-specific recombination in mice. PNAS 93: 10887–10890. DOI: https://doi.org/10.1073/pnas.93.20.10887, PMID: 8855277

13. Floor S, van Staveren WC, Larsimont D, Dumont JE, Maenhaut C. 2011. Cancer cells in epithelial-to-mesenchymal transition and tumor-propagating-cancer stem cells: distinct, overlapping or same populations. Oncogene 30: 4609–4621. DOI: https://doi.org/10.1038/onc.2011.184, PMID: 21643013

14. Grigorenko BL, Nemukhin AV, Shadrina MS, Topol IA, Burt SK. 2007. Mechanisms of guanosine triphosphate hydrolysis by Ras and Ras-GAP proteins as rationalized by ab initio QM/MM simulations. Proteins 66: 456–466. DOI: https://doi.org/10.1002/prot.21228, PMID: 17094109

15. Gebregiworgis T, Kano Y, St-Germain J, Radulovich N, Udaskin ML, Mentes A, Huang R, Poon BPK, He W, Valencia-Sama I, Robinson CM, Huestis M, Miao J, Yeh JJ, Zhang ZY, Irwin MS, Lee JE, Tsao MS, Raught B, Marshall CB, et al. 2021. The Q61H mutation decouples KRAS from upstream regulation and renders cancer cells resistant to SHP2 inhibitors. Nature Communications 12: 6274. DOI: https://doi.org/10.1038/s41467-021-26526-y, PMID:34725361

16. Hahn WC, Dessain SK, Brooks MW, King JE, Elenbaas B, Sabatini DM, DeCaprio JA, Weinberg RA. 2002. Enumeration of the simian virus 40 early region elements necessary for human cell transformation. Molecular and Cellular Biology 22: 2111–2123. DOI: https://doi.org/10.1128/mcb.22.7.2111-2123.2002, PMID: 11884599

17. Hobbs GA, Baker NM, Miermont AM, Thurman RD, Pierobon M, Tran TH, Anderson AO, Waters AM, Diehl JN, Papke B, Hodge RG, Klomp JE, Goodwin CM, DeLiberty JM, Wang J, Ng RWS, Gautam P, Bryant KL, Esposito D, Campbell SL, et al. 2020. Atypical KRAS(G12R) mutant is impaired in PI3K signaling and macropinocytosis in pancreatic cancer. Cancer Discovery 10: 104–123. DOI: https://doi.org/10.1158/2159-8290.CD-19-1006, PMID: 31649109

18. Huber W, Carey VJ, Gentleman R, Anders S, Carlson M, Carvalho BS, Bravo HC, Davis S, Gatto L, Girke T, Gottardo R, Hahne F, Hansen KD, Irizarry RA, Lawrence M, Love MI, MacDonald J, Obenchain V, Oleś AK, Pagès H, et al., 2015. Orchestrating high-throughput genomic analysis with Bioconductor. Nature Methods 12: 115–121. DOI: https://doi.org/10.1038/nmeth.3252, PMID: 25633503

19. Jackson EL, Willis N, Mercer K, Bronson RT, Crowley D, Montoya R, Jacks T, Tuveson DA. 2001. Analysis of lung tumor initiation and progression using conditional expression of oncogenic K-ras. Genes & Development 15: 3243–3248. DOI: https://doi.org/10.1101/gad.943001, PMID: 11751630

20. Kersey PJ, Staines DM, Lawson D, Kulesha E, Derwent P, Humphrey JC, Hughes DS, Keenan S, Kerhornou A, Koscielny G, Langridge N, McDowall MD, Megy K, Maheswari U, Nuhn M, Paulini M, Pedro H, Toneva I, Wilson D, Yates A, et al., 2012. Ensembl Genomes: an integrative resource for genome-scale data from non-vertebrate species. Nucleic Acids Research 40: D91–D97. DOI: https://doi.org/10.1093/nar/gkr895, PMID: 22067447

21. Klaeger S, Heinzlmeir S, Wilhelm M, Polzer H, Vick B, Koenig PA, Reinecke M, Ruprecht B, Petzoldt S, Meng C, Zecha J, Reiter K, Qiao H, Helm D, Koch H, Schoof M, Canevari G, Casale E, Depaolini SR, Feuchtinger A, et al., 2017. The target landscape of clinical kinase drugs. Science 358: eaan4368. DOI: https://doi.org/10.1126/science.aan4368, PMID: 29191878

22. Kong G, Chang YI, You X, Ranheim EA, Zhou Y, Burd CE, Zhang J. 2016. The ability of endogenous Nras oncogenes to initiate leukemia is codon-dependent. Leukemia 30: 1935–1938. DOI: https://doi.org/10.1038/leu.2016.89, PMID: 27109513

23. Kotting C, Kallenbach A, Suveyzdis Y, Wittinghofer A, Gerwert K. 2008. The GAP arginine finger movement into the catalytic site of Ras increases the activation entropy. PNAS 105: 6260–6265. DOI: https://doi.org/10.1073/pnas.0712095105, PMID: 18434546

24. Lake D, Correa SA, Muller J. 2016. Negative feedback regulation of the ERK1/2 MAPK pathway. Cellular and Molecular Life Sciences, 73: 4397–4413. DOI: https://doi.org/10.1007/s00018-016-2297-8, PMID: 27342992

25. Lampson BL, Pershing NL, Prinz JA, Lacsina JR, Marzluff WF, Nicchitta CV, MacAlpine DM, Counter CM. 2013. Rare codons regulate KRas oncogenesis. Current Biology 23: 70–75. DOI: https://doi.org/10.1016/j.cub.2012.11.031, PMID: 23246410

26. Li S, Balmain A, Counter CM. 2018. A model for RAS mutation patterns in cancers: finding the sweet spot. Nature Reviews Cancer 18: 767–777. DOI: https://doi.org/10.1038/s41568-018-0076-6, PMID: 32286309

27. Liu F, Yang X, Geng M, Huang M. 2018. Targeting ERK, an Achilles’ Heel of the MAPK pathway, in cancer therapy. Acta Pharmaceutica Sinica B, 8: 552–562. DOI: https://doi.org/10.1016/j.apsb.2018.01.008, PMID: 30109180

28. Liu P, Jenkins NA, Copeland NG. 2003. A highly efficient recombineering-based method for generating conditional knockout mutations. Genome Research 13: 476–484. DOI: https://doi.org/10.1101/gr.749203, PMID: 12618378

29. Love MI, Huber W, Anders S. 2014. Moderated estimation of fold change and dispersion for RNA-seq data with DESeq2. Genome Biology 15: 550. DOI: https://doi.org/10.1186/s13059-014-0550-8, PMID: 25516281

30. Lu S, Jang H, Nussinov R, Zhang J. 2016. The Structural Basis of Oncogenic Mutations G12, G13 and Q61 in Small GTPase K-Ras4B. Scientific Reports 6: 21949. DOI: https://doi.org/10.1038/srep21949, PMID 26902995

31. Mani SA, Guo W, Liao MJ, Eaton EN, Ayyanan A, Zhou AY, Brooks M, Reinhard F, Zhang CC, Shipitsin M, Campbell LL, Polyak K, Brisken C, Yang J, Weinberg RA. 2008. The epithelial-mesenchymal transition generates cells with properties of stem cells. Cell 133: 704–715. DOI: https://doi.org/10.1016/j.cell.2008.03.027, PMID: 18485877

32. Marjanovic ND, Hofree M, Chan JE, Canner D, Wu K, Trakala M, Hartmann GG, Smith OC, Kim JY, Evans KV, Hudson A, Ashenberg O, Porter CBM, Bejnood A, Subramanian A, Pitter K, Yan Y, Delorey T, Phillips DR, Shah N, et al., 2020. Emergence of a high-plasticity cell state during lung cancer evolution. Cancer Cell 38: 229–246. DOI: https://doi.org/10.1016/j.ccell.2020.06.012, PMID: 32707077

33. Martin M. 2011. Cutadapt removes adapter sequences from high-throughput sequencing reads. EMBnet.journal 17: 10–12. DOI :https://doi.org/10.14806/ej.17.1.200.

34. Matkar SS, Durham A, Brice A, Wang TC, Rustgi AK, Hua X. 2011. Systemic activation of K-ras rapidly induces gastric hyperplasia and metaplasia in mice. American Journal of Cancer Research, 1: 432–445. PMID:21761008

35. Matys V, Fricke E, Geffers R, Gössling E, Haubrock M, Hehl R, Hornischer K, Karas D, Kel AE, Kel-Margoulis OV, Kloos DU, Land S, Lewicki-Potapov B, Michael H, Münch R, Reuter I, Rotert S, Saxel H, Scheer M, Thiele S, et al., 2003. TRANSFAC: transcriptional regulation, from patterns to profiles. Nucleic Acids Research 31: 374–378. DOI: https://doi.org/10.1093/nar/gkg108, PMID: 12520026

36. Matys V, Kel-Margoulis OV, Fricke E, Liebich I, Land S, Barre-Dirrie A, Reuter I, Chekmenev D, Krull M, Hornischer K, Voss N, Stegmaier P, Lewicki-Potapov B, Saxel H, Kel AE, Wingender E. 2006. TRANSFAC and its module TRANSCompel: transcriptional gene regulation in eukaryotes. Nucleic Acids Research 34: D108–D110. DOI: https://doi.org/10.1093/nar/gkj143, PMID: 16381825

37. Mootha VK, Lindgren CM, Eriksson KF, Subramanian A, Sihag S, Lehar J, Puigserver P, Carlsson E, Ridderstråle M, Laurila E, Houstis N, Daly MJ, Patterson N, Mesirov JP, Golub TR, Tamayo P, Spiegelman B, Lander ES, Hirschhorn JN, Altshuler D, et al., 2003. PGC-1alpha-responsive genes involved in oxidative phosphorylation are coordinately downregulated in human diabetes. Nature Genetics 34: 267–273. DOI: https://doi.org/10.1038/ng1180, PMID: 12808457

38. Muñoz-Maldonado C, Zimmer Y, Medová M. 2019. A comparative analysis of individual RAS mutations in cancer biology. Frontiers in Oncology 9: 1088. DOI: https://doi.org/10.3389/fonc.2019.01088, PMID: 31681616

39. Nakamura Y, Gojobori T, Ikemura T. 2000. Codon usage tabulated from international DNA sequence databases: status for the year 2000. Nucleic Acids Research 28: 292. DOI: https://doi.org/10.1093/nar/28.1.292, PMID: 10592250

40. O’Hayer KM, Counter CM. 2006. A genetically defined normal human somatic cell system to study Ras oncogenesis in vivo and in vitro. Methods in Enzymology, 407: 637–647. DOI: https://doi.org/10.1016/S0076-6879(05)07050-3, PMID: 16757358

41. Parikh N, Shuck RL, Nguyen TA, Herron A, Donehower LA. 2012. Mouse tissues that undergo neoplastic progression after K-Ras activation are distinguished by nuclear translocation of phospho-Erk1/2 and robust tumor suppressor responses. Molecular Cancer Research 10: 845–855. DOI: https://doi.org/10.1158/1541-7786.MCR-12-0089, PMID: 22532587

42. Parker JA, Volmar AY, Pavlopoulos S, Mattos C. 2018. K-Ras populates conformational states differently from its isoform H-Ras and oncogenic mutant K-RasG12D. Structure 26: 810–820 e814. DOI: https://doi.org/10.1016/j.str.2018.03.018, PMID: 29706533

43. Pershing NL, Lampson BL, Belsky JA, Kaltenbrun E, MacAlpine DM, Counter CM. 2015. Rare codons capacitate Kras-driven de novo tumorigenesis. Journal of Clinical Investigations 125: 222–233. DOI: https://doi.org/10.1172/JCI77627, PMID: 25437878

44. Peterson J, Li S, Kaltenbrun E, Erdogan O, Counter CM. 2020. Expression of transgenes enriched in rare codons is enhanced by the MAPK pathway. Scientific Reports 10: 22166. DOI: https://doi.org/10.1038/s41598-020-78453-5, PMID: 33335127

45. Poulin EJ, Bera AK, Lu J, Lin YJ, Strasser SD, Paulo JA, Huang TQ, Morales C, Yan W, Cook J, Nowak JA, Brubaker DK, Joughin BA, Johnson CW, DeStefanis RA, Ghazi PC, Gondi S, Wales TE, Iacob RE, Bogdanova L, Gierut JJ, et al. 2019. Tissue-specific oncogenic activity of KRAS(A146T). Cancer Discovery 9: 738–755. DOI: https://doi.org/0.1158/2159-8290.CD-18-1220, PMID: 30952657

46. Prior IA, Hood FE, Hartley JL. 2020. The frequency of Ras mutations in cancer. Cancer Research 80: 2969–2974. DOI: https://doi.org/10.1158/0008-5472.CAN-19-3682, PMID: 32209560

47. Rabara D, Tran TH, Dharmaiah S, Stephens RM, McCormick F, Simanshu DK, Holderfield M. 2019. KRAS G13D sensitivity to neurofibromin-mediated GTP hydrolysis. PNAS 116: 22122–22131. DOI: https://doi.org/10.1073/pnas.1908353116, PMID 31611389

48. Ray KC, Bell KM, Yan J, Gu G, Chung CH, Washington MK, Means AL. 2011. Epithelial tissues have varying degrees of susceptibility to Kras(G12D)-initiated tumorigenesis in a mouse model. PLoS One 6: e16786. DOI: https://doi.org/10.1371/journal.pone.0016786, PMID: 21311774

49. Rouillard AD, Gundersen GW, Fernandez NF, Wang Z, Monteiro CD, McDermott MG, Ma’ayan A. 2016. The harmonizome: a collection of processed datasets gathered to serve and mine knowledge about genes and proteins. Database 2016: baw100. DOI: https://doi.org/10.1093/database/baw100, PMID: 27374120

50. Scheffzek K, Ahmadian MR, Kabsch W, Wiesmüller L, Lautwein A, Schmitz F, Wittinghofer A. 1997. The Ras-RasGAP complex: structural basis for GTPase activation and its loss in oncogenic Ras mutants. Science 277: 333–338. DOI: https://doi.org/10.1126/science.277.5324.333, PMID: 9219684

51. Schneider CA, Rasband WS, Eliceiri KW. 2012. NIH Image to ImageJ: 25 years of image analysis. Nature Methods 9: 671–675. DOI: https://doi.org/10.1038/nmeth.2089, PMID: 22930834

52. Serrano M, Lin AW, McCurrach ME, Beach D, Lowe SW. 1997. Oncogenic ras provokes premature cell senescence associated with accumulation of p53 and p16INK4a. Cell 88: 593–602. DOI: https://doi.org/10.1016/s0092-8674(00)81902-9, PMID: 9054499

53. Sharp PM, Li WH. 1987. The codon Adaptation Index--a measure of directional synonymous codon usage bias, and its potential applications. Nucleic Acids Research 15: 1281–1295. DOI: https://doi.org/10.1093/nar/15.3.1281, PMID: 3547335

54. Simanshu DK, Nissley DV, McCormick F. 2017. RAS proteins and their regulators in human disease. Cell 170: 17–33. DOI: https://doi.org/10.1016/j.cell.2017.06.009, PMID: 28666118

55. Singh K, Pruski M, Bland R, Younes M, Guha S, Thosani N, Maitra A, Cash BD, McAllister F, Logsdon CD, Chang JT, Bailey-Lundberg JM. 2020. Kras mutation rate precisely orchestrates ductal derived pancreatic intraepithelial neoplasia and pancreatic cancer. Laboratory Investigations, 101: 177–192. DOI: https://doi.org/10.1038/s41374-020-00490-5, PMID: 33009500

56. Smith MJ, Neel BG, Ikura M. 2013. NMR-based functional profiling of RASopathies and oncogenic RAS mutations. PNAS 110: 4574–4579. DOI: https://doi.org/10.1073/pnas.1218173110, PMID: 23487764

57. Sutherland KD, Song JY, Kwon MC, Proost N, Zevenhoven J, Berns A. 2014. Multiple cells-of-origin of mutant K-Ras-induced mouse lung adenocarcinoma. PNAS 111: 4952–4957. DOI: https://doi.org/10.1073/pnas.1319963111, PMID: 24586047

58. Tata PR, Chow RD, Saladi SV, Tata A, Konkimalla A, Bara A, Montoro D, Hariri LP, Shih AR, Mino-Kenudson M, Mou H, Kimura S, Ellisen LW, Rajagopal J. 2018. Developmental history provides a roadmap for the emergence of tumor plasticity. Developmental Cell 44: 679–693. DOI: https://doi.org/10.1016/j.devcel.2018.02.024, PMID: 29587142

59. To MD, Wong CE, Karnezis AN, Del Rosario R, Di Lauro R, Balmain A. 2008. Kras regulatory elements and exon 4A determine mutation specificity in lung cancer. Nature Genetics, 40: 1240–1244. DOI: https://doi.org/10.1038/ng.211, PMID 18758463

60. Ventura A, Kirsch DG, McLaughlin ME, Tuveson DA, Grimm J, Lintault L, Newman J, Reczek EE, Weissleder R, Jacks, T. 2007. Restoration of p53 function leads to tumour regression in vivo. Nature 445: 661–665. DOI: https://doi.org/10.1038/nature05541, PMID: 17251921

61. Wang L, Jin Q, Lee JE, Su IH, Ge K. 2010. Histone H3K27 methyltransferase Ezh2 represses Wnt genes to facilitate adipogenesis. PNAS 107: 7317–7322. DOI: https://doi.org/10.1073/pnas.1000031107, PMID: 20368440

62. Winters IP, Chiou SH, Paulk NK, McFarland CD, Lalgudi PV, Ma RK, Lisowski L, Connolly AJ, Petrov DA, Kay MA, Winslow MM. 2017. Multiplexed in vivo homology-directed repair and tumor barcoding enables parallel quantification of Kras variant oncogenicity. Nature Communications 8: 2053. DOI: https://doi.org/10.1038/s41467-017-01519-y, PMID: 29233960

63. Wong JC, Perez-Mancera PA, Huang TQ, Kim J, Grego-Bessa J, Del Pilar Alzamora M, Kogan SC, Sharir A, Keefe SH, Morales CE, Schanze D, Castel P, Hirose K, Huang GN, Zenker M, Sheppard D, Klein OD, Tuveson DA, Braun BS, Shannon K. 2020. KrasP34R and KrasT58I mutations induce distinct RASopathy phenotypes in mice. JCI Insight 5: e140495. DOI: https://doi.org/10.1172/jci.insight.140495, PMID: 32990679

64. Xu X, Rock JR, Lu Y, Futtner C, Schwab B, Guinney J, Hogan BL, Onaitis MW. 2012. Evidence for type II cells as cells of origin of K-Ras-induced distal lung adenocarcinoma. PNAS 109: 4910–4915. DOI: https://doi.org/10.1073/pnas.1112499109, PMID 22411819

65. Zafra MP, Parsons MJ, Kim J, Alonso-Curbelo D, Goswami S, Schatoff EM, Han T, Katti A, Fernandez MTC, Wilkinson JE, Piskounova E, Dow LE. 2020. An in vivo Kras allelic series reveals distinct phenotypes of common oncogenic variants. Cancer Discovery 10: 1654–1671. DOI: https://doi.org/10.1158/2159-8290.CD-20-0442, PMID: 32792368

